# Zebrafish models of human-duplicated *SRGAP2* reveal novel functions in microglia and visual system development

**DOI:** 10.1101/2024.09.11.612570

**Authors:** José M. Uribe-Salazar, Gulhan Kaya, KaeChandra Weyenberg, Brittany Radke, Keiko Hino, Daniela C. Soto, Jia-Lin Shiu, Wenzhu Zhang, Cole Ingamells, Nicholas K. Haghani, Emily Xu, Joseph Rosas, Sergi Simó, Joel Miesfeld, Tom Glaser, Scott C. Baraban, Li-En Jao, Megan Y. Dennis

## Abstract

The expansion of the human *SRGAP2* family, resulting in a human-specific paralog *SRGAP2C,* likely contributed to altered evolutionary brain features. The introduction of *SRGAP2C* in mouse models is associated with changes in cortical neuronal migration, axon guidance, synaptogenesis, and sensory-task performance. Truncated SRGAP2C heterodimerizes with the full-length ancestral gene product SRGAP2A and antagonizes its functions. However, the significance of *SRGAP2* duplication beyond neocortex development has not been elucidated due to the embryonic lethality of complete *Srgap2* knockout in mice. Using zebrafish, we show that *srgap2* knockout results in viable offspring and that these larvae phenocopy “humanized” *SRGAP2C* larvae, including altered morphometric features (i.e., reduced body length and inter-eye distance) and differential expression of synapse-, axonogenesis-, and vision-related genes. Through single-cell transcriptome analysis, we demonstrate a skewed balance of excitatory and inhibitory neurons that likely contribute to increased susceptibility to seizures displayed by *Srgap2* mutant larvae, a phenotype resembling *SRGAP2* loss-of-function in a child with early infantile epileptic encephalopathy. Single-cell data also shows strong endogenous expression of *srgap2* in microglia with mutants exhibiting altered membrane dynamics and likely delayed maturation of microglial cells. Microglia cells expressing *srgap2* were also detected in the developing eye together with altered expression of genes related to axonogenesis in mutant retinal cells. Consistent with the perturbed gene expression in the retina, we found that *SRGAP2* mutant larvae exhibited increased sensitivity to broad and fine visual cues. Finally, comparing the transcriptomes of relevant cell types between human (+*SRGAP2C*) and non-human primates (–*SRGAP2C*) revealed significant overlaps of gene alterations with mutant cells in our zebrafish models; this suggests that *SRGAP2C* plays a similar role altering microglia and the visual system in modern humans. Together, our functional characterization of conserved ortholog Srgap2 and human SRGAP2C in zebrafish uncovered novel gene functions and highlights the strength of cross-species analysis in understanding the development of human-specific features.

**Abstract (short):** *SRGAP2C* has been implicated in contributing to altered brain features in the evolution of humans. However, the significance of *SRGAP2* duplication beyond neocortex development has not been elucidated due to the embryonic lethality of complete *Srgap2* knockout in mice. Using zebrafish, we show that *srgap2* knockout results in viable offspring that phenocopy “humanized” *SRGAP2C* larvae. Morphometric, behavioral, and transcriptome analyses collectively suggest *srgap2* impacts axonal guidance, synaptogenesis, and seizure susceptibility. Beyond neurons, *Srgap2* functions in controlling membrane dynamics and maturation of microglial cells, possibly leading to altered axonogenesis in the developing retina and increased sensitivity to broad and fine visual cues. Comparing relevant transcriptomes between human and nonhuman primates suggests that *SRGAP2C* similarly impacts microglia and vision in modern humans. Our functional characterization of conserved ortholog Srgap2 and human SRGAP2C in zebrafish uncovered novel gene functions and highlights the strength of cross-species analysis in understanding the development of human-specific features.

## Introduction

Genetic factors contributing to phenotypic differences between humans and non-human primates remain largely undiscovered ^1,2^. However, gene expansions ^3,4^ have been suggested as an important driver of primate species divergence ^5–13^, with mammalian and organoid models recapitulating hallmark features of human brain development, including altered synaptogenesis, corticogenesis, and gyrification ^14–20^. One of the most well-studied human duplicated genes is the Slit-Robo Rho GTPase-activating protein 2 (*SRGAP2*) ^14,18,21–25^. Multiple *SRGAP2* paralogs arose over the last ∼3.4 million years, resulting in a conserved ancestral full-length *SRGAP2* and three truncated human-specific paralogs (*SRGAP2B*, *SRGAP2C*, and a likely nonfunctional *SRGAP2D* ^23^), all located on human chromosome 1 (Figure 1A, top). The human ancestral *SRGAP2A* encodes a protein with F-BAR, RhoGAP, and SH3 domains, while *SRGAP2B* and *SRGAP2C* encode only F-BAR domains ^26^. SRGAP2 forms homodimers with itself through the F-BAR domain; SRGAP2B and SRGAP2C dimerize with the F-BAR domain of SRGAP2A, leading to degradation of the resulting heterodimer via the proteasome pathway ^21,24^. SRGAP proteins modulate cytoskeleton dynamics and promote membrane deformation when dimerized impacting vital cellular processes such as motility, polarity, and morphogenesis ^26^ (Figure 1A, bottom).

**Figure 1.**
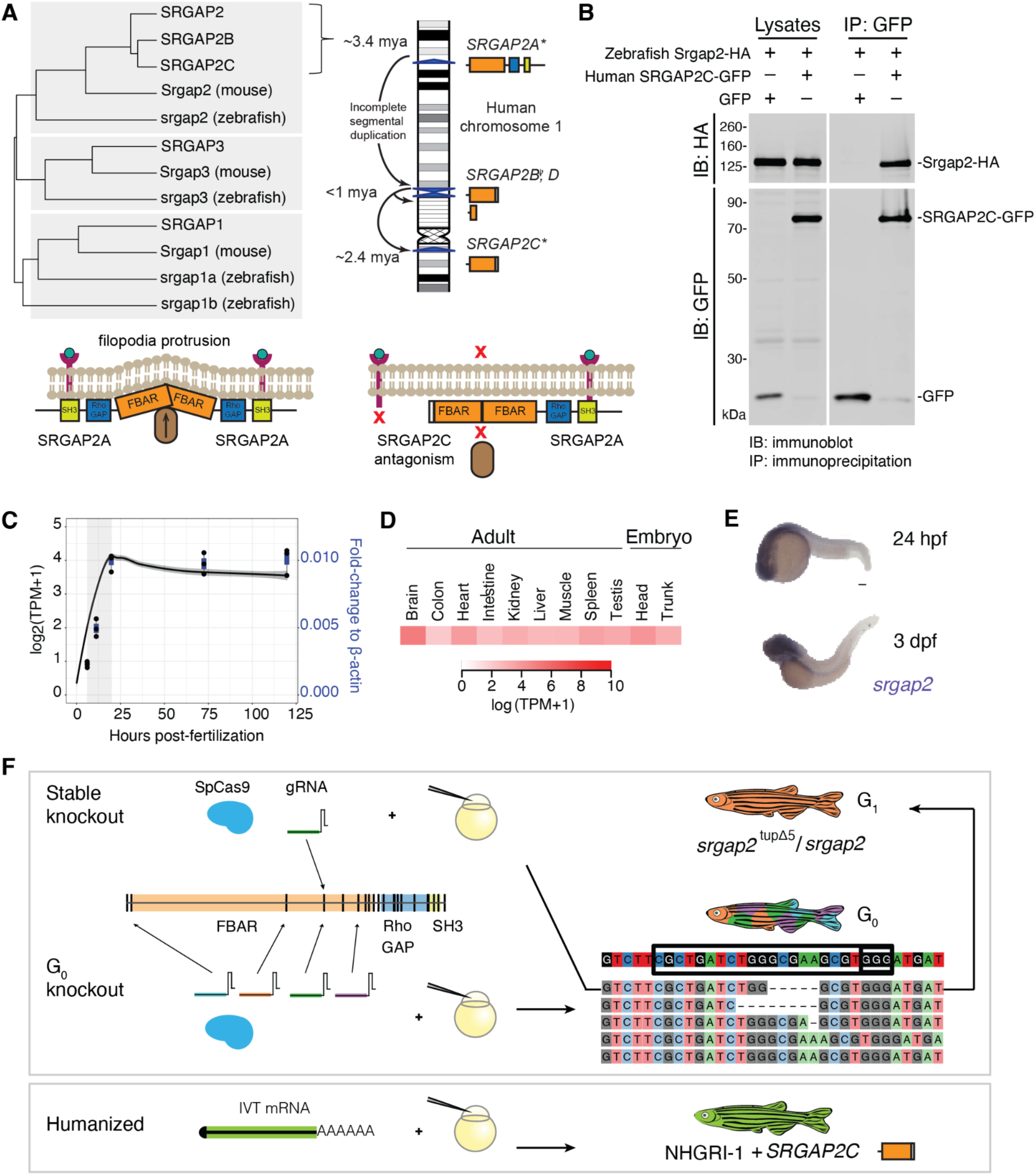
Functional analysis of *srgap2* in the developing zebrafish. **(A)** Top left, phylogenetic tree of human, mice, and zebrafish SRGAP proteins based on their full length amino acid sequence using the Unweighted Pair Group method with Arithmetic Mean method. Top right, schematic of inferred *SRGAP2* gene family evolutionary history across human chromosome 1 ^25^. Bottom, cartoon summarizing the results of previous studies, showing that SRGAP2 functions after homodimerization in concert with F-actin (brown oval) to dictate cell membrane dynamics (bottom left) or heterodimerize with SRGAP2C producing no functional product (bottom right). (**B)** Co-immunoprecipitation of human-specific SRGAP2C and zebrafish Srgap2 in HEK293T cells showed interaction between these proteins. (**C)** Temporal expression of *srgap2* in the developing embryo according to publicly available RNA-seq data ^31^ (black line represents the best fit line with the standard error in dark gray) and normalized quantitative RT-PCR data from whole-embryo RNA collected at 6, 10, 24, 72, and 120 hpf (blue boxes, each dot represents a biological replicate). The light-gray box represents a critical stage in zebrafish neurogenesis between 6 and 24 hpf ^32^. (**D)** *srgap2* expression in embryonic (24 hpf) and adult (>12 months old) tissues from a published RNA-seq dataset ^34^. (**E)** Spatial endogenous expression of *srgap2* at 24 hpf and 3 dpf via *in situ* hybridization shown in blue. Scale bar 100 µm. **(F)** Illustration of the approaches to creating knockout *srgap2* zebrafish. Top, a stable knockout line was generated by injecting SpCas9 coupled with one gRNA targeting exon 4. Middle, G_0_ knockouts were generated by co-injecting SpCas9 coupled with four gRNAs targeting early exons. Bottom, humanized larvae were created by injecting *in vitro* transcribed *SRGAP2C* mRNA at the one-cell stage.

Mouse *Srgap2* has important functions in synapse maturation and connectivity via interactions with Homer, Gephyrin, and Rac1, the known regulators of both excitatory and inhibitory synapse maturation ^18,24^. Mouse models expressing human *SRGAP2C* consistently phenocopy *Srgap2* knockdown and conditional knockout mice, showing conserved functional antagonism between the human and mouse proteins. These models exhibit increases in the rate of neuronal migration, neurite outgrowth, and density of dendritic spines, as well as neoteny in the spine maturation process ^18,21,24^. Expressing *SRGAP2C* also increases long-range synaptic connectivity in mouse cortical pyramidal neurons and enhanced cortical processing abilities in the whisker-based texture-discrimination tests ^22^. Together, these studies support the contribution of *SRGAP2C* to the emergence of unique neuronal features and cognitive capacities in humans.

Human-specific traits exist beyond the neocortex, including alterations in the development of musculoskeletal features and the eye ^27^. The broad expression of *SRGAP2* paralogs in human cells and tissues ^6^ suggests it may have functions outside of neurons. However, the embryonic lethality of complete *Srgap2* loss-of-function in mouse models ^28^ allows limited assessment of its global functions. Here, we report that zebrafish *srgap2* “knockout” models result in viable offspring, allowing us to characterize *SRGAP2* developmental functions beyond the neocortex. We compared s*rgap2* knockouts with *SRGAP2C*-expressing “humanized” larvae in a range of morphological, gene expression, cellular, molecular, and behavioral assays. We observed concordant effects in *srgap2* knockout and *SRGAP2C*-humanized larvae across all assays, demonstrating that human-specific SRGAP2C antagonizes zebrafish Srgap2 functions and verifying previous known functions of *SRGAP2* as an axon/synapse regulator. We found zebrafish mutants exhibit increased susceptibility to seizures, strengthening previous findings in a human patient ^29^, present evidence that *SRGAP2* is a conserved core gene in microglia function across vertebrates that alters membrane dynamics and delays maturation, and propose a never-before-reported role of *SRGAP2* in the developing eye that impacts vision. Combined, our zebrafish models support previous studies in mice and expand on exciting new functions of *SRGAP2C*, opening new lines of inquiry related to microglia and the retina in the evolution of human-specific traits.

## Results

### Genomic and transcriptional conservation of the zebrafish *srgap2* ortholog

The current zebrafish genome (GRCz11/danRer11) carries a single *srgap2* ortholog encoding F-BAR, RhoGAP, and SH3 domains. Human full-length SRGAP2 and zebrafish Srgap2 proteins share 73.8% amino acid identity, placing them phylogenetically closer to each other than to other members of the SRGAP protein family (Figure 1A). The F-BAR domain of zebrafish Srgap2 shares 87.9% amino acid identity with that of human SRGAP2. Computationally-predicted ^30^ probabilities of interaction between SRGAP2C and zebrafish Srgap2 were comparable to those between SRGAP2C and mouse Srgap2 ^18,21,24^ (Table S1). We experimentally confirmed heterodimer formation between zebrafish Srgap2 and human-specific SRGAP2C by performing co-immunoprecipitation in HEK293T cells (Figure 1B).

Published whole-embryo RNA sequencing (RNA-seq)^31^ showed that expression of *srgap2* continues to increase after fertilization, plateaus after around 16 hours post fertilization (hpf), and persists thereafter; we confirmed this expression pattern with quantitative RT-PCR (Figure 1C). Thus, the initiation of *srgap2* expression coincides with critical neurogenesis periods, including the formation of post-mitotic neurons in the neural plate after gastrulation ^32^ between 5.25 and 10 hpf ^33^. Tissue-specific RNA-seq data from embryos (24 hpf) and adults (12 months old) ^34^ showed high *srgap2* expression in the embryonic head and adult brain with lower expression in viscera (e.g., heart, spleen, and kidney; Figure 1D). To validate these results, we performed whole-mount *in situ* hybridization and observed *srgap2* expression mainly in the developing central nervous system at 24 hpf and 3 days post fertilization (dpf) (Figure 1E). Overall, *srgap2* expression is spatiotemporally regulated during a critical period of early neurodevelopment in the zebrafish embryo and remains high in the adult brain ^34^. These results suggest that zebrafish can serve as a suitable model to test *SRGAP2* paralog functions during neural development.

### *SRGAP2C* humanized larvae phenocopy *srgap2* knockout models

We evaluated *SRGAP2* function during development using two different zebrafish knockout models (Figure 1F). First, we generated a stable *srgap2* knockout line carrying a 5-bp deletion in exon 4 using CRISPR-Cas9 mutagenesis (*srgap2^tupΔ5^*, Table S2). While stable mutant zebrafish lines are classically used to test gene function ^35,36^, we also created mosaic embryos carrying a mixture of *srgap2* knockout alleles by injecting ribonucleoproteins containing SpCas9 coupled with four different guide RNAs (gRNAs) targeting early exons (termed “G_0_ knockouts”) ^37–40^. Evaluation of control (wild type (WT) or scrambled-injected) and mutant larvae revealed significantly decreased *srgap2* mRNA abundance at 5 dpf in both knockout models (average relative reductions versus WT: Het= 23.8%, Hom= 59.7%, G_0_ knockouts= 55.6%; Figure S1A). We observed no detectable off-target mutations in knockout larvae from either approach at the most probable sites predicted using CIRCLE-Seq and CRISPRScan (Table S3), suggesting that any observed phenotypes were due to the loss of Srgap2 function.

Homozygous (Hom) or heterozygous (Het) siblings produced from crossing *srgap2* knockout (Het) parents showed no difference in mortality at 5 dpf from WT siblings (survival curve test: χ^2^= 2.96, df= 2, *p*-value= 0.228, n=148, WT= 19%, Het= 53%, Hom= 28%). G_0_-knockouts also showed similar mortality to scrambled gRNA controls (χ^2^= 0.3, df= 1, *p*-value= 0.6, n G_0_-knockouts= 347, n controls= 260). However, we observed significant reductions in the length of the body axis (∼4.4–7.6%) and distance between the eyes (∼1.5–4.7%) of all *srgap2* knockout larvae (Het, Hom, and G_0_) versus controls (Figure 2A). No significant effects on head-trunk angle, a feature typically used to estimate developmental timing in early zebrafish larvae ^33^, nor head area were observed, allowing us to rule out developmental delay (Figure 2A, S1B). Given the similarity of morphological features in both stable and mosaic knockout models, we primarily focused on phenotypes produced in G_0_ knockout mutants moving forward.

**Figure 2.**
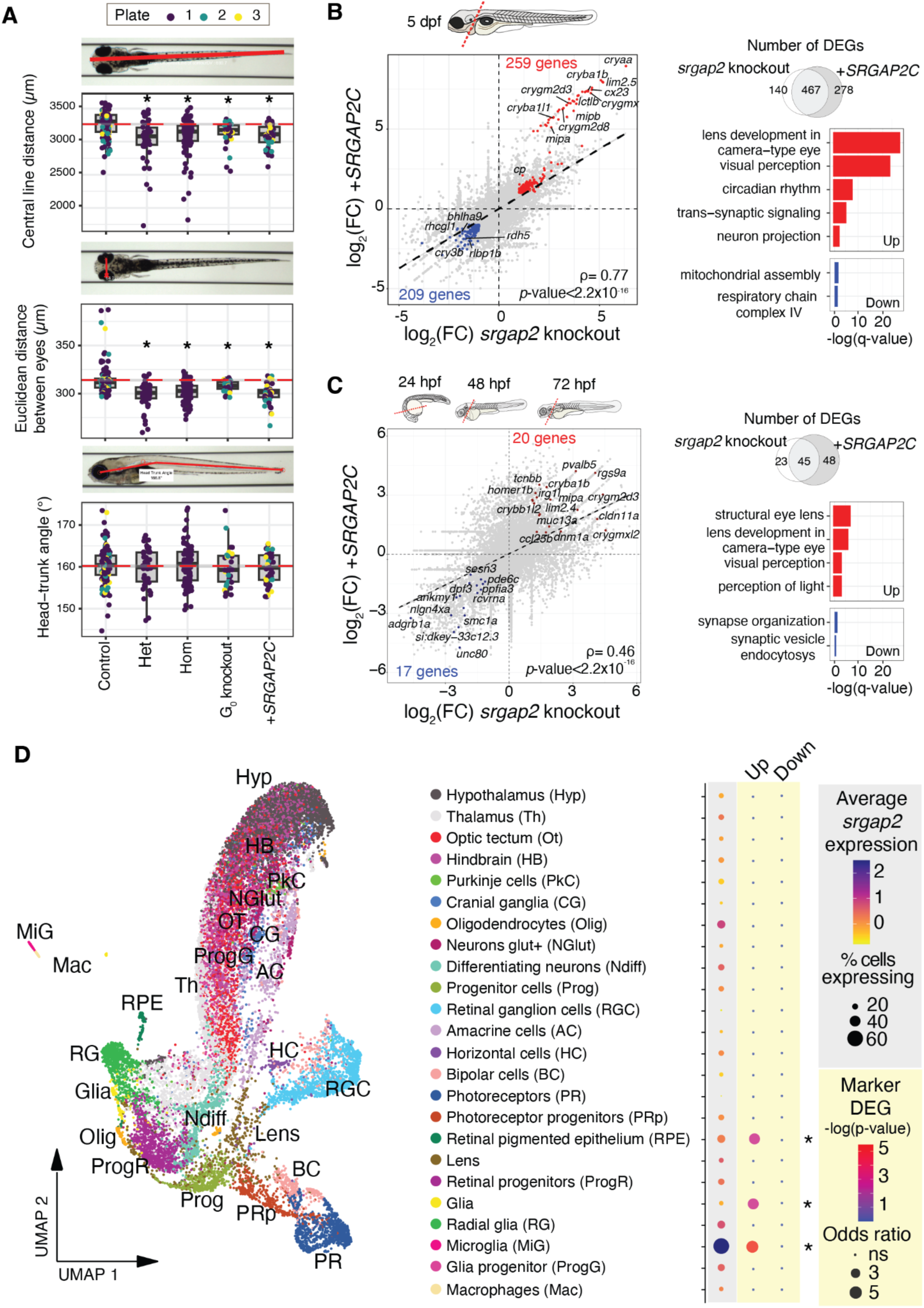
Developmental and cellular phenotypes of diverse zebrafish models of *SRGAP2*. (**A)** Measurements of central line distance (ANOVA: F_(4, 321)_= 12.84, genotype effects *p*-value= 1.04x10^-9^, FDR-adjusted *p*-values Het= 4.40x10^-7^, Hom= 6.29x10^-7^, Pooled= 0.015, *SRGAP2C*= 1.36x10^-4^), Euclidean distance between the eyes (ANOVA: F_(4,321)_= 23.49, genotype effects *p*-value= 4.72x10^-17^, Dunnett’s test FDR-adjusted *p*-values: Het= 6.77x10^-11^, Hom= 4.69x10^-10^, Pooled= 0.05, *SRGAP2C*= 2.19x10^-9^), and head angle (ANOVA: F_(4,315)_= 0.49, genotype effects *p*-value=0.746) in 5 dpf larvae from stable *srgap2* knockout (Het n= 43, Hom n= 86), G_0_ knockouts (n= 34), *SRGAP2C*-injected (n= 44), and control larvae (n= 124). Dots represent an imaged larva with the color indicating the imaging plate (a co-variable included in the statistical analyses). The red dotted line corresponds to the mean value for the control group. Representative images of each measurement are included on the top of each plot. (**B)** Correlation of the fold change (FC) between *srgap2* G_0_-knockouts and *SRGAP2C*-injected larvae at 5 dpf, with common DEGs highlighted (red= upregulated (FC > 2), blue= downregulated (FC < -2)). Top representative GO terms enriched in common DEGs between *srgap2* G_0_-knockouts and *SRGAP2C*-injected larvae (complete results in Table S5). Color of the bar represents the direction of the genes (red= commonly upregulated, blue= commonly downregulated). (**C)** Correlation of the FC between *srgap2* G_0_-knockouts and *SRGAP2C*-injected larvae across development using data from 24, 48, and 72 hpf larvae, with common DEGs highlighted, complete results can be found in Tables S7, S8. (**D)** Clustering of the 28,687 profiled cells colored as 24 cell types based on the expression of gene markers. Expression of *srgap2* across cell types (left side, shaded in gray), with the size of the circle representing the percentage of cells in that cluster expressing *srgap2* and the color of the circle representing the average scaled expression in the cluster. Enrichment test for the overlap between marker genes for each cell type and the differentially expressed genes at 3 dpf from bulk RNA-seq data (right side), with the size of the circle representing the odds ratio for the enrichment and the color of the circle the -log(BH-adjusted *p-*value) of the Fisher’s exact test. Asterisks indicate an FDR-adjusted *p*-value < 0.05.

Next, we generated a *SRGAP2C* humanized model by microinjecting *in vitro* transcribed mRNA into one-cell stage embryos (Figure 1F). This produced transient and ubiquitous presence of *SRGAP2C* transcripts in the developing zebrafish up to 72 hpf (Figure S1A), coinciding with peak endogenous *srgap2* expression (starting at 16 hpf; Figure 1C), with protein likely persisting for longer. *SRGAP2C-*humanized larvae developed normally with no increased mortality (survival curve test: χ^2^= 0.8, df= 1, *p*-value= 0.4, *SRGAP2C*-injected= 422, eGFP-mRNA-injected controls= 308). They exhibited significant changes in overall body length and distance between the eyes (∼5.7% reduction in body length and ∼4.2% reduction in distance between the eyes Figure 2A), similar to the phenotypes observed in the knockout models. Thus, introducing human SRGAP2C antagonized endogenous zebrafish Srgap2 function in developing zebrafish larvae, similar to what has been observed in the mouse models ^14,18,21,24^.

### Transcriptomes reveal developmental impacts upon perturbation of Srgap2 function

Given that knocking out *srgap2* and expressing human *SRGAP2C* generated similar developmental phenotypes (Figure 2A), we reasoned that a common set of molecular processes were perturbed under these two experimental conditions. To test this, we performed RNA-seq of dissected heads from G_0_ knockouts and *SRGAP2C*-injected embryos/larvae across early developmental stages and performed differential expression analysis versus respective controls. From this, we observed high correlation of expression changes (Figures 2B and C, Note S1) and significant enrichment of shared differentially expressed genes (DEGs) between the knockout and humanized models (e.g., 467 shared genes at 5 dpf, Fisher’s exact test odds ratio= 378.3, *p*-value < 2.2x10^-16^, Table S4). We next assessed enriched gene ontology (GO) of shared DEGs between *srgap2* G_0_ knockout and *SRGAP2C* humanized larvae (which we collectively term “*SRGAP2* mutants”) to understand molecular changes in these models.

We found that shared upregulated genes across all developmental time points were related to lens and visual system development in the *SRGAP2* mutant models (Tables S4–S8). Upregulated genes unique in older mutant larvae (5 dpf) were related to neurodevelopment (mainly neuronal projections and synapse organization) and circadian rhythm, and downregulated genes involved mitochondrial cytochrome c oxidase assembly (Figures 2A). Mitochondrial dysfunction is associated with reduced height ^41^, consistent with the reduced body axis observed in *SRGAP2* mutant larvae. Young mutant embryos (1, 2, 3 dpf) exhibited downregulation of genes related to synapse organization relative to the controls, suggesting delayed synaptic maturation. In particular, *ppfia3*, a regulator of presynapse assembly ^42^, was found significantly upregulated in 5 dpf mutant larvae while downregulated in mutant embryos (≤ 3 dpf). These results align with the neoteny of synaptogenesis observed in *Srgap2* knockdown or *SRGAP2C*-expressing mouse embryos ^18^. In summary, identifying the common pathways affected in the *srgap2* knockout and *SRGAP2C*-humanized zebrafish models provide evidence of the critical role that the genes play in the development of the visual system and neurodevelopment.

To narrow in on the cell types driving expression changes, we performed transcriptomic profiling (SPLiT-seq ^43^) of 28,687 single cells isolated from 3 dpf zebrafish larval brains of *SRGAP2* mutants and controls (Table S9). Using expression patterns of marker genes ^44,45^, we classified 24 cell types and found broad endogenous *srgap2* expression across neuron-containing clusters, with highest expression in microglia (Figure 2D, Tables S10 and S11). We observed a significant enrichment of upregulated DEGs we associated with *SRGAP2* mutants through bulk RNA-seq analysis (Figure 2C) in markers for retinal pigmented epithelium (RPE), glia, and microglia cells (Table S12).

### Synaptic alterations in *SRGAP2* zebrafish models

Based on the broad neural expression of *srgap2*, we next performed differential pseudo-bulk differential expression analysis using 11,450 neuronal cells. Shared upregulated genes between *SRGAP2* mutant models (n=14) related to neuron projection guidance (Figure 3A), including *ephb2*, which is implicated in promoting/directing axon guidance across the brain midline ^46,47^. Shared downregulated genes (n=21) were enriched for synaptic signaling functions, concordant with bulk RNA-seq results (Figure 3A, Tables S13 and S14). With respect to neuronal subtypes, markers for forebrain (comprising the telencephalon and orthologous to the mammalian neocortex ^48,49^), midbrain (composed of optic tectum, the visual processing center in the zebrafish brain ^50^), and differentiating neurons were enriched in upregulated genes; while hindbrain and the broad neuron category were enriched for downregulated genes (Figure 3A, Table S15, BH-adjusted Fisher’s exact tests *p*-values < 0.05).

**Figure 3.**
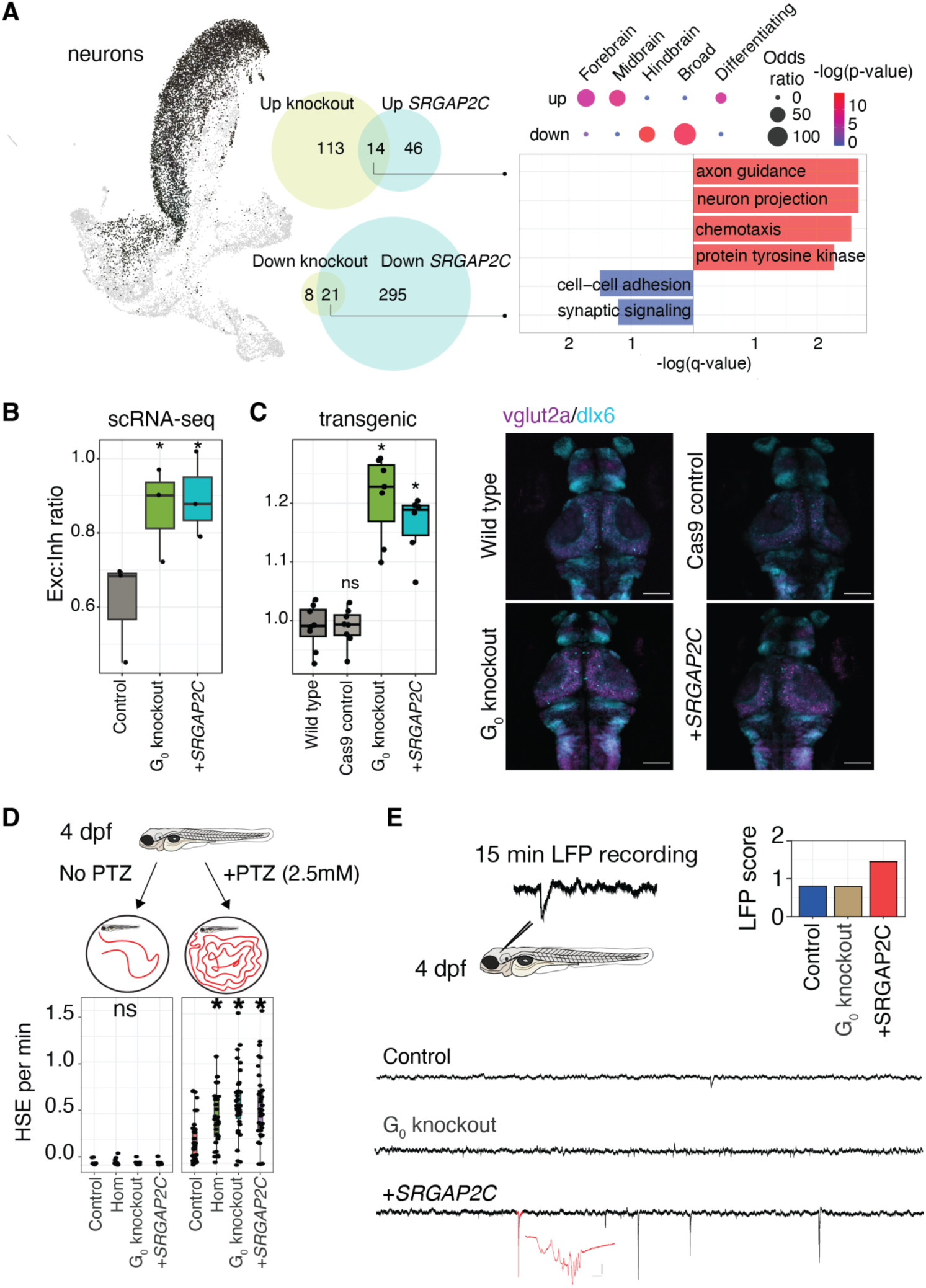
Neuronal alterations in *SRGAP2* mutants. (**A)** Neuronal clusters (hypothalamus, thalamus, optic tectum, hindbrain, Purkinje cells, and neurons rich in glutamate receptors) selected to perform a differential gene expression test was performed to DEGs in the *SRGAP2* mutants compared to the control group. Bar plot represents the top GO terms overrepresented in the 14 commonly upregulated genes (complete results in Table S14). (**B)** Ratio of cells classified as excitatory (*vglut2^+^*) to inhibitory (*gad1b^+^*) between the *srgap2* G_0_-knockouts, *SRGAP2C*-injected, and controls (*srgap2* G_0_ knockouts: 0.78±0.15, *p*-value= 0.031; *SRGAP2C*-injected: 0.82±0.09, *p*-value= 0.017, controls= 0.57±0.13; t-tests versus controls). (**C)** Ratio of excitatory (*vglut2*:DsRed*+*) to inhibitory (*dlx6*:GFP) cell area quantified from images of 3 dpf *srgap2* G_0_-knockout, *SRGAP2C*-injected, SpCas9 control injected, and uninjected wild type larvae (G_0_ knockout: Exc:Inh ratio=1.21±0.07, *p*-value=3.0x10^-4^, *SRGAP2C*: Exc:Inh ratio= 1.16±0.05, *p*-value= 7.0x10^-4^, SpCas9-injected controls Exc:Inh ratio= 0.98±0.03, p-value= 0.959; Mann-Whitney U-tests *p*-values vs wild-type controls). Images include representative samples per group, scale bars 100 µm. (**D)** High-speed events (HSE, >28 mm/s) identified in 15 min recordings of 4 dpf larvae (*srgap2* knockouts (stable Hom_parent_ and G_0_), *SRGAP2C*-injected, and SpCas9-injected controls, n= 36 larvae per group) with and without PTZ. Frequency of HSE per min were compared to controls (0 mM PTZ: ANOVA *p*-value for genotypic effect= 0.415, average HSE/min= 0.006±0.02, no significant differences between groups; 2.5 mM PTZ: ANOVA genotype effect *p*-value= 1.1x10^-6^, Hom_parent_= 0.010, G_0_-knockouts= 2.2x10^-6^, *SRGAP2C*-injected= 3.90x10^-5^). **(E)** Local field potential (LFP) recordings in the optic tectum of 4 dpf larvae (G_0_-knockouts, *SRGAP2C*-injected, and SpCas9-injected controls, n=21-30 per group) were obtained and scored by two independent researchers. Representative traces per group are shown. Asterisks in graphs represent a *p*-value below 0.05 for the comparison against the control group. ns= not significant.

Given findings of altered synaptic signaling/organization in mutant zebrafish (Figure 2B) and the role of *SRGAP2* paralogs in regulating synapses in mice ^18^, we narrowed in on excitatory (Exc; *slc17a6b*/*vglut2*) and inhibitory (Inh; *gad1b*) neuronal subtypes in our scRNA-seq data ^44,45^. Comparing relative abundances across models showed that both *srgap2* knockouts and *SRGAP2C*-injected larvae exhibited a ∼20% increase in the Exc:Inh ratio (Figure 3B). Quantifying co-labeled GABAergic (Tg[*dlx6a*:GFP] ^51^) and glutamatergic (Tg[*vglut2a:*DsRed] ^52^) neurons validated these results, with a ∼29% increase in the Exc:Inh ratio relative to uninjected wild-type and control-injected larvae (Figure 3C). The ratio measured in our controls matched that from previous studies using the same transgenic lines of the same age ^53^ (wild-type controls Exc:Inh ratio= 0.98±0.04).

A skew towards excitatory versus inhibitory neuronal balance is associated with seizures, as has been reported in several zebrafish epilepsy models ^54^. We therefore assessed chemically-induced seizure-like behaviors in control and *SRGAP2* mutant models ^55^. We counted high-speed movement events (HSE, >28 mm/s) in 4 dpf larvae exposed to either a low concentration of pentylenetetrazol (PTZ, 2.5 mM) or to E3 media (control). While HSE were rare in non-PTZ-treated larvae with no difference in frequency between genotypic groups (average HSE/min= 0.006±0.02; Figure 3D), the addition of PTZ significantly increased the frequency of HSE on average by 0.31±0.08 min^-1^ in *srgap2* knockouts and *SRGAP2C*-humanized larvae compared to controls. Next, we detected spontaneous electrographic seizures by recording local field potentials (LFP) ^55^. *SRGAP2C* larvae demonstrated ictal-like Type II electrical events, classifying them as epileptic (n= 21, LFP score= 1.45, Figure 3E), while control (n= 22) and *srgap2* G_0_-knockouts (n= 30) did not exhibit any events. Strikingly, *SRGAP2C* larvae showed LFP scores in the range observed in zebrafish models of well-established epilepsy-associated genes (e.g., *SCN1A*, *STXBP1*) ^55^. Overall, our results point to a role for SRGAP2 and its human-specific paralog *SRGAP2C* in maintaining neuronal E:I balance and potentially contributing to seizure susceptibility.

### *SRGAP2* is a conserved microglial gene that inhibits cell ramifications

While *SRGAP2* functions are well-characterized in neurons, we observed the highest expression of *srgap2* in microglia (Figure 3D). This observation is concordant with a previous study implicating *SRGAP2* as a “core” microglia gene with high conservation across human, macaque, marmoset, sheep, rat, mouse, hamster, and zebrafish ^56^. When comparing the transcriptomes of microglia cells from *srgap2* knockout and *SRGAP2C*-expressing zebrafish models versus their respective controls, we found that shared upregulated genes (n=38) were implicated in cell migration and shared downregulated genes (n=65) were related to actin-mediated filopodia processes (Figure 4A, Tables S16 & S17). These results align with the ability of SRGAP2 to induce cell projections in concert with F-actin in a variety of systems ^57,58^. Since microglia also develop complex cell ramifications, we hypothesized that their cell-membrane dynamics were also modulated by Srgap2 activity.

**Figure 4.**
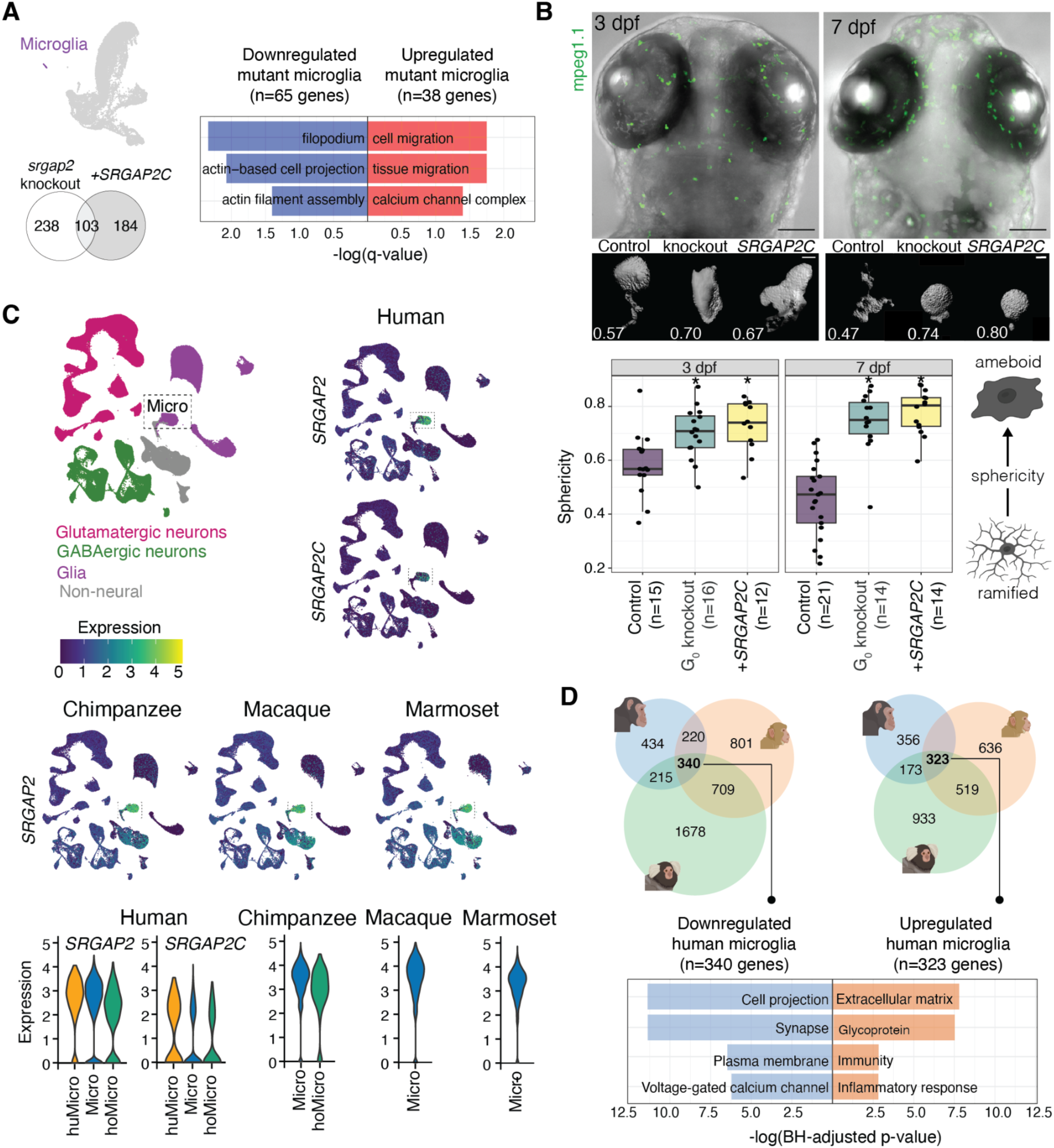
Cross-species conservation of *SRGAP2* as a microglial gene. (**A)** Top GO terms with significant overrepresentation in genes upregulated (red) or downregulated (blue) in microglial cells from *SRGAP2* mutants from Figure 2D. (**B)** Sphericity values for individual microglial cells (mpeg1.1^+^) at 3 and 7 dpf in *srgap2* knockouts, *SRGAP2C*-injected, and scrambled gRNA-injected controls. Each dot represents a single microglial cell (average of 4-5 cells per larvae from 3-4 larvae per genotype per timepoint were obtained). Representative images for the median sphericity value of larvae at 3 and 7 dpf for each genotype are included below the graph (scale bars: top images= 100µm, bottom images= 5 µm). Asterisks denote a Tukey *post*-hoc p-value < 0.05. 3dpf: *srgap2* G_0_ knockouts: 0.70±0.09, *p*-value= 0.0085; *SRGAP2C*-injected: 0.73±0.09, *p*-value= 0.0021, controls: 0.58±0.12; 7dpf: *srgap2* G_0_ knockouts: 0.74±0.11, *p*-value < 2.2x10^-16^; *SRGAP2C*-injected: 0.78±0.08, *p*-value < 2.2x10^-16^, controls: 0.46±0.13. (**C)** Evaluation of 610,596 prefrontal cortex cells from human, chimpanzee, macaque, and marmoset (human: 171,997, chimpanzee: 158,099, macaque: 131,032, marmoset: 149,468) showing the levels of *SRGAP2* and *SRGAP2C* expression across species, highlighting the microglial cluster with a dotted square. Micro: microglia. Expression of *SRGAP2* and *SRGAP2C* in microglial subtypes across species with subtypes ordered from highest expression left to right. huMicro: human-specific microglia, hoMicro: Hominidae-specific microglia. **(D)** Microglial cells from human, chimpanzee, macaque, and marmoset (human: 8,819 cells, chimpanzee: 6,000 cells, macaque: 9,000 cells, marmoset: 7,099 cells) from the prefrontal cortex and middle temporal gyrus were used to identify common DEGs between human and non-human primates, finding 340 common upregulated and 323 common downregulated genes. Top GO terms with significant overrepresentation in common DEGs are included.

To test this, we characterized microglia in *srgap2* G_0_ knockouts and humanized *SRGAP2C* models. While there was no difference in microglia abundance between genotypes ^59^ (Figure S2), we observed significantly reduced ramifications (quantified as increased sphericity) for both knockout and humanized larvae compared to controls at both 3 and 7 dpf using a transgenic line labeling macrophages (Tg[*mpeg1.1*:GFP], Figure 4B) ^60^. By these developmental time points, macrophages are generally accepted to be microglia (or their precursors) when localized in the brain/retina of zebrafish ^61^. The microglia in control larvae continued to acquire more ramified morphologies from 3 to 7 dpf as they matured (t-test of 3 vs 7 dpf: t= 2.97, *p*-value= 0.0055, Figure 4B), concordant with previous reports ^62^. Microglia in both *srgap2* mutant models retained similar sphericity at both timepoints (t-tests per mutant genotype *p*-values > 0.05), suggesting arrested maturation. However, we cannot rule out increased microglia activation, which also involves morphological changes from a ramified “resting” state to more ameboid-like active shapes ^63,64^. This interpretation is supported, in part, by the upregulation of known microglial activation markers (*hsp90aa1.1* and *zfp36l2*) observed in our *SRGAP2* mutants at 5 dpf ^65^ from bulk RNA-seq results (Figure 2B).

To ask if SRGAP2/C might contribute to human-specific microglia membrane dynamics, we re-analyzed published single-cell transcriptomes of 610,596 prefrontal cortex cells from human, chimpanzee, macaque, and marmoset ^66^. In line with its conserved “core” characterization ^56^*, SRGAP2* exhibited highest expression in the microglia clusters in all primates (Figure 4C), including human- and Hominidae-specific microglia subclusters (Figure 4D, Note S2, Table S18). *SRGAP2C* expression was also high in all human microglia subtypes, albeit slightly lower compared to *SRGAP2*. Taking a pseudo-bulk approach analogous to the zebrafish analysis (Figure 3A), we compared differential expression of human (+*SRGAP2C*) versus chimpanzee, macaque, or marmoset (-*SRGAP2C* “controls”) microglia. Human DEGs were consistent with reduced microglia ramifications, including downregulation of genes associated with cell projection and the plasma membrane (Table S19). We also observed the upregulation of genes implicated in extracellular matrix and inflammatory response, both features of migrating microglia in an ameboid state. Comparing DEGs between human/primate and zebrafish *SRGAP2* mutants revealed significant overlap (10 genes, Fisher’s test odds ratio= 2.77, *p*-value= 0.0046). Thus, the alterations of microglial cell shape observed in our zebrafish *SRGAP2C* “humanized” models were recapitulated in human-specific biological processes that occur in microglial cells.

### Visual system alterations in *SRGAP2* zebrafish models

The most striking molecular change in *SRGAP2* mutant zebrafish was the upregulation of genes related to lens development and visual perception (Figure 2B & C). Performing RNA *in situ* hybridization (ISH) of the developing zebrafish eye, we found predominant endogenous *srgap2* expression in the optic nerve (ON), RPE, and along the retinal ganglion cell layer (GCL) at 3 dpf (Figure 5A). While scRNA-seq data showed strong expression of *srgap2* and enrichment of differential marker genes in RPE cells, we found little to no *srgap2* expression in retinal ganglion cells (RGCs) comprising the GCL (Figure 2D). Instead, *srgap2* ISH likely marks microglia that have migrated into the retina, with strongest expression evident at the interface between the lens and the neural retina.

**Figure 5.**
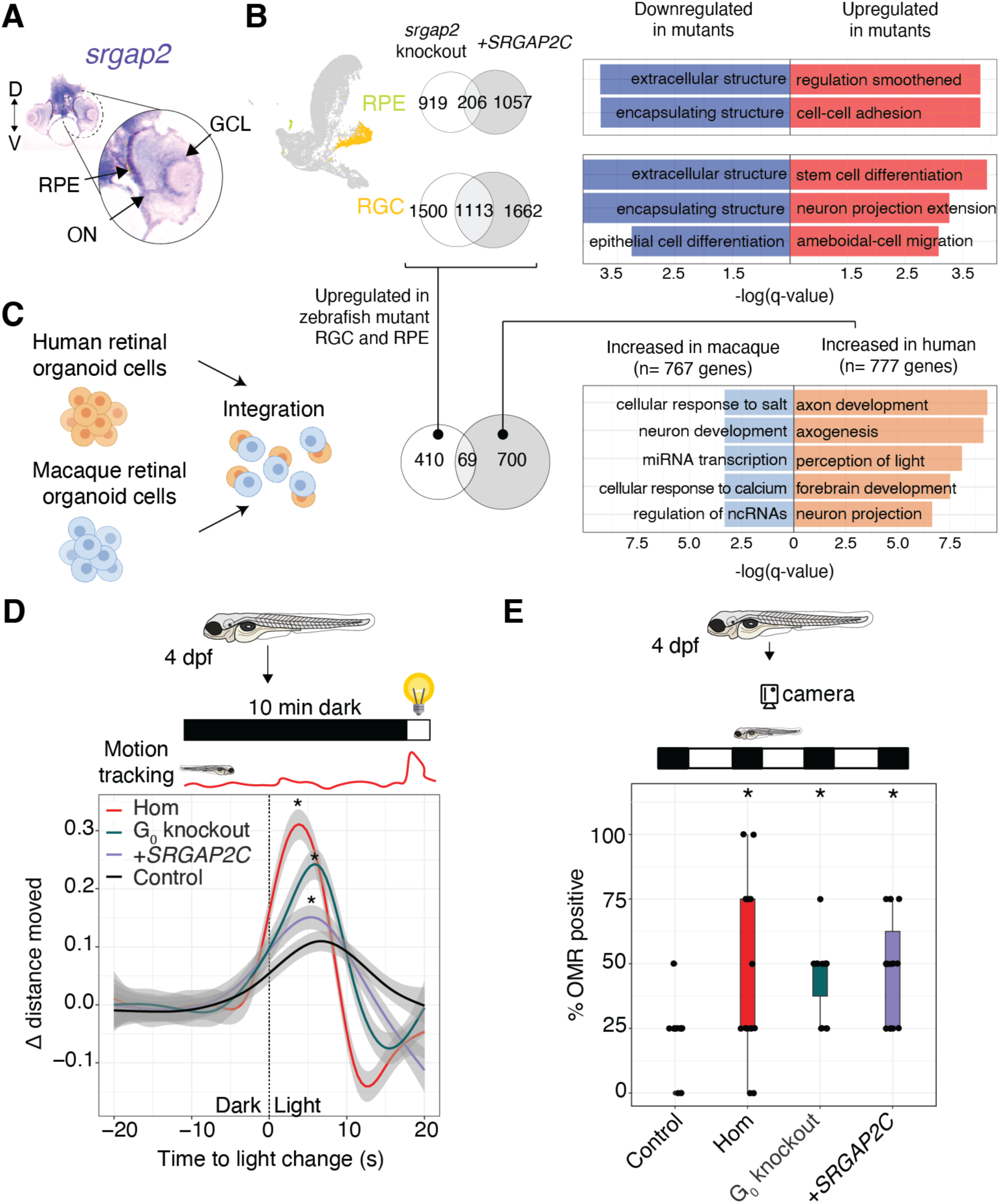
*SRGAP2* impacts the retina. (**A)** Section of a 3 dpf NHGRI-1 larva staining *srgap2* expression via *in situ* hybridization, labeling predominantly the optic nerve (ON), retinal pigmented epithelium (RPE), and the ganglion cell layer (GCL). D: dorsal, V: ventral. (**B)** Retinal ganglion cells (RGCs) were selected and a differential gene expression performed between *SRGAP2*-mutants (*srgap2* knockouts and *SRGAP2C*-injected) versus controls, identifying 60 upregulated genes and 84 downregulated genes, with their top overrepresented GO terms included in bar plots. **(C)** Human and macaque cells from retinal organoids (43,857 human and 19,894 macaque) were integrated to identify genes with increased expression in either species, with their top overrepresented GO terms included in bar plots (complete results in Tables S22 and S23). (**D)** Motion response to changes in light were assessed in 4 dpf *srgap2* knockouts (Hom_parent_ and G_0_-knockouts), *SRGAP2C*-injected, and SpCas9-scrambled gRNA-coupled control larvae using a 10 min acclimation period followed by an abrupt light change. Plot includes trend lines for change in distance moved observed in each evaluated group (n= 24 per group, standard error for each line included as a shaded gray), which were different between all groups compared to controls (Kolmogorov-Smirnov tests *p*-values: Hom_parent_= 9.16x10^-11^, G_0_-knockouts= 5.93x10^-8^, *SRGAP2C*-injected= 1.11x10^-12^). (**E)** Optomotor responses were evaluated in 4 dpf larvae using an optimized protocol^73^ that quantifies the percentage of larvae relative to moving stripes. Boxplot includes the percentage of OMR-positive larvae (aligned to the visual stimulus) in *srgap2* knockouts (Hom_parent_ and G_0_-knockouts) and *SRGAP2C*-injected, which was higher compared to controls (Dunn’s Benjamini-Hochberg adjusted *p*-values: Hom_parent_= 0.0113, G_0_-knockouts= 0.0040, *SRGAP2C*-injected= 0.0040). Asterisks denote a *p*-value below 0.05.

To understand biological impacts within the retina, we identified DEGs across RGCs and RPE cells in *SRGAP2* mutants versus controls. RGCs were enriched for shared upregulated genes related to stem-cell differentiation, neuron-projection extension, and amoeboid-type cell migration (Figure 5B, Tables S20 & S21). Shared upregulated genes in RPE were also associated with cell-cell adhesion as well as negative regulation of the smoothened pathway, which mediates response to Hedgehog signaling ^67^. Shared downregulation of genes important in extracellular structures (e.g., matrix metalloproteinases, laminin, and collagen gene families) was observed in both RGCs and the RPE. Connecting our findings to the developing human retina (organoids ^68,69^ and post mortem ^70,71^), transcriptomic data from human (+*SRGAP2C*) versus rhesus macaque (-*SRGAP2C* “controls”) also show upregulation of similar pathways related to axon development and neuron projections. Again, we observed a significant overlap in common DEGs between cells from human retina and *SRGAP2* zebrafish mutant RGC/RPE (69 genes, Fisher’s test odds ratio= 6.23, *p*-value< 2.2x10^-16^; Tables S22-S25 and Note S3). Importantly, the eyes of *srgap2* knockout and *SRGAP2C*-humanized zebrafish developed normally with the formation of all major cell types by 5 dpf, indicating that these changes in gene expression did not affect gross aspects of eye development (Figure S3). Together, these results point to unexplored human-specific eye development features facilitated by SRGAP2C—related to membrane dynamics impacting axonogenesis—altering retinal connectivity that is fundamental for visual information processing ^72^.

To test if the observed differences in gene expression patterns are associated with altered vision, we leveraged natural zebrafish larval behavior that react to abrupt changes in light intensity with increased swimming activity ^74,75^. Using motion tracking, we observed a significant increase in response (reaction time and movement) to light stimulus in *srgap2* mutants (knockouts and *SRGAP2C*-injected) compared to controls (Figure 5D) at 4 dpf, showing increased sensitivity to light changes. Considering our models exhibited increased susceptibility to seizures, which could evoke similar responses, we also characterized more refined visual cues. The optomotor response (OMR) measures the instinctive behavior of free-swimming zebrafish larvae wherein they align their body axis in the same direction as contrasting visual stimuli, such as moving stripes. This helps freshwater fish swim upstream ^73,76,77^. We found that a larger percentage of 4 dpf *srgap2* knockouts (Hom_parent_ and G_0_) and *SRGAP2C*-humanized mutants showed OMR-positive positioning compared to the control group (n per group= 15, Figure 5E). Together, these results suggest that reduction in Srgap2 activity—either through genetic knockouts or expression of human *SRGAP2C*—impacts the function of retinal microglia and possibly contributes to altered neuronal connectivity in the developing eye, leading to more sensitive neuronal responses to visual cues.

## Discussion

*SRGAP2* is a well-studied human-specific duplicated gene with a wealth of gain- and loss-of-function studies in diverse cell culture and mouse models. Its documented functions include regulating neuronal migration, synaptogenesis, and long-range connectivity in the central nervous system ^14,18,22,24^. However, because of the embryonic lethality of the *Srgap2* knockout in mouse models, any roles beyond the neocortex are still largely unexplored. Here, we present new functional analyses of *SRGAP2* in zebrafish, where viable knockout mutants allow detailed screening of developmental phenotypes at an organismal level. We observed an overall concordance in developmental phenotypes between *srgap2* knockouts and *SRGAP2C*-injected zebrafish larvae, similar to previous mouse studies where temporal expression of truncated *SRGAP2C* mirrored *Srgap2*-knockdown/knockout alleles ^14,18,24^. For example, both *SRGAP2* knockout and humanized models consistently exhibited shorter body length, a phenotype not reported previously. This could be driven by altered mitochondrial functions as suggested by bulk RNA-seq analysis (Figure 2B), or by perturbation to migration-dependent processes such as muscle guidance and body patterning that are influenced by the Slit-Robo pathway ^78,79^. No heterozygous loss-of-function variants have been discovered in ancestral *SRGAP2* across hundreds of thousands of healthy humans to date ^80^, indicating strong selective constraint (gnomAD pLI=0.87); moving forward, it will be interesting to compare mutant phenotypes with those of human patients carrying gene-impacting mutations.

Bulk transcriptomic analyses of mutant zebrafish—ranging from 24 hpf embryos to 5 dpf larvae— revealed alterations to known molecular functions, suggesting increased axonogenesis in *SRGAP2* mutants consistent with the gene’s well-characterized role in axonal guidance via the Slit-Robo pathway ^14^. Downregulation of genes related to synaptogenesis in early developmental embryos (24 hpf–3 dpf, Figure 2C) is concordant with neoteny of synaptogenesis in *SRGAP2* mouse models reminiscent of human brain development ^18,24^. The single-cell transcriptomes allowed us to further narrow in on altered neuronal functions (Figure 2D). For example, mutants exhibited skewed Exc:Inh balance of neurons that manifested as increased susceptibility to chemically-induced seizures (Figure 3C). *SRGAP2C*-expressing larvae also presented spontaneous, unprovoked, electrographic seizures not observed in our G_0_ knockout mutant. Differences in phenotypic severity between the knockout and humanized models might be explained by genetic compensation due to nonsense-mediated decay in the knockout mutant ^81^. Transcriptome data of *SRGAP2* mutant neurons provided additional clues to possible mechanisms underlying the observed phenotypes; for example, we observed significantly reduced expression of the *GRIN2A* ortholog (*grin2ab*, Table S13), encoding glutamate [NMDA] receptor subunit epsilon-1, with loss-of-function variants implicated in epileptic aphasia in humans ^82^. These results are largely consistent with a clinical report of early infantile epileptic encephalopathy in a human child carrying a reciprocal translocation disrupting *SRGAP2* ^29^, providing evidence that mutations of this gene may contribute to epilepsy. We note that the embryonic lethality of *Srgap2* knockout mice has impeded similar evaluations in mammalian models to date.

Hallmark studies have shown that *Srgap2* loss-of-function or *SRGAP2C* expression leads to reduced filopodia in COS7 cells and fewer branching processes in mouse cortical neurons ^14^ altering neuronal migration *in vivo* ^21^. Similarly, we found that mutant zebrafish microglia exhibit reduced ramifications versus controls, also evident in transcriptomes (reduced expression of filopodia and actin-based cell projections-related genes and increased expression of cell migration genes). The mutant microglia also maintained an ameboid-like spherical shape through development time (3 to 7 dpf; Figure 4B) instead of the expected increased ramifications observed in a typically-developing zebrafish larva ^62^. This ameboid-like shape is indicative of either “active” or immature microglia. While we cannot rule out that mutant microglia were more activated, we propose that microglia exhibited developmental delay like that observed in synaptic spine maturation in mice ^21^. Indeed, a recent preprint ^83^ showed similar microglia neoteny in *SRGAP2C* mouse and human cell models. Interestingly, human adult microglia also express *SRGAP2* paralogs and exhibit similar transcriptome differences with nonhuman primates as *SRGAP2C*-humanized zebrafish microglia do with controls. Most overlapping DEGs function in actin-cytoskeleton dynamics (down) and cell-cell interactions (up). This provides molecular evidence of altered membrane dynamics of human microglia compared with other primates, consistent with the reduced ramifications observed for adult human microglia compared with macaque and marmoset imaged from post-mortem brain samples ^56^.

The most striking results produced by our transcriptomic analysis implicates vision development in *SRGAP2* mutants, a function never-before reported in genetic models of *SRGAP2*. Crystallins were amongst the highest upregulated genes found at 5 dpf (Figure 2B). While these genes are typically associated with lens development, we observed no gross morphological defects in the lenses of stable homozygous knockout larvae or adults (data not included). We did find *srgap2* to be highly expressed in axonal-rich regions of the zebrafish eyes (ON and retinal GCL), in line with *Srgap2* expression observed in mouse GCL ^84^. Interestingly, upregulation of crystallin genes has also been reported in the retinas of *Srgap2*^+/-^ adult mice ^28^. Alpha-crystallins are small heat-shock proteins that have been associated with axonal elongation ^85^ and regeneration ^86^. Similarly, we axonogenesis genes were upregulated in both SRGAP2 mutant zebrafish and human retinal organoids, when compared to a nonhuman primate (rhesus macaque). Connecting possible axonal guidance changes with vision ^87^, we tested visual-motor responses of zebrafish larvae to abrupt light-dark changes or moving contrast stimuli ^74,88^ and consistently show that *srgap2* knockout and *SRGAP2C*-expressing larvae have an increased response to visual cues, suggestive of enhanced visual processing.

Given the presence of *srgap2*-expressing microglia in the developing zebrafish eye, we propose a model where predominantly-amoeboid mutant microglia plays a role in retinal axon extension. Microglia are resident macrophages in the brain that migrate into the central nervous system early in development, influencing wide-ranging developmental processes such as synaptogenesis and pruning, neurogenesis, and axonogenesis ^89,90^. The eye is among the first regions to be colonized by microglia, at ∼26–30 hpf in zebrafish ^59^, with preferential localization to differentiating cells in the retina GCL ^61^ (also evident in our Tg[*mpeg1.1*:GFP] lines at 3 dpf, Figure 4B). *SRGAP2*-mutant microglia, in their immature and potentially activated state, could play a role in increased clearance of dead/apoptotic cells or pruning axons/synapses leading to altered retinal connectivity and improved visual processing. Further, beyond impacts in the eye, it is plausible that microglia mediate other brain phenotypes observed in *SRGAP2* mutant zebrafish. This has recently been proposed for changes in synaptic development of cortical pyramidal neurons observed in a microglia-specific *Srgap2* conditional knockout mouse model ^83^. While we have yet to directly connect *SRGAP2*-related microglia functions to the observed changes in Exc:Inh neuronal balance of our mutant zebrafish, studies have found that microglial activation induces increased frequency of excitatory synaptic events ^91^. Microglia are also associated with pro- and anti-epileptic activity due to their various roles in brain homeostasis and neuroinflammation ^92^ suggesting possible connections with seizures detected in our *SRGAP2* mutants. Moving forward, generation of microglia-specific *SRGAP2* zebrafish models will allow us to delineate microglia functions in retina and brain development.

While our studies using zebrafish have allowed us to query novel *SRGAP2* functions at an organismal level, they also present some limitations. “Humanizing” larvae by injection of *SRGAP2C* mRNA at the single-cell stage introduces the gene ubiquitously, possibly contributing to off-target phenotypes. However, all published studies to date show SRGAP2C functions solely by antagonizing *srgap2,* suggesting that the truncated human-specific paralog would be non-functional on its own. A strength of this approach is that SRGAP2C-driven antagonism potentially produces more severe phenotypes, as it avoids the genetic compensation that can occur in knockout models ^81^. This might explain differences in fold-change of DEGs between *srgap2* knockout and humanized models (Figure 2B), in particular across vision-related genes (Note S1). Nevertheless, to avoid possible confounding factors, our conservative transcriptome analysis considered only DEGs observed in both knockout and humanized *SRGAP2* models. Further, because SRGAP2C was transiently introduced, we only characterized phenotypes in zebrafish larvae up to 7 dpf, limiting the scope of our study to early developmental traits. Finally, the structure of the zebrafish forebrain, which lacks a neocortex, limits analysis of certain processes specific to mammals, such as subtle circuit changes between cortical regions observed in *SRGAP2* mouse models ^22^. Regardless, conservation at cellular and molecular levels has successfully enabled zebrafish models of neurodevelopmental conditions impacting the cortex, such as autism and intellectual disability, across hundreds of genes ^93–98^.

In summary, we have leveraged the advantages of viable zebrafish *SRGAP2* mutant models to investigate its functional roles. Our findings are concordant with previous reports implicating *SRGAP2* in neurological phenotypes and reveal novel functions in microglia and the developing eye. Combined, these results provide new hypotheses regarding *SRGAP2C*-driven changes to microglia function and axonogenesis in the brain and retina unique to humans, as well as improvements in visual perception, opening many avenues to test in cross-species comparisons moving forward.

## Methods

### Zebrafish lines and husbandry

NHGRI-1 wild type zebrafish lines ^99^ were maintained using standard protocols ^100^. Animals were maintained in a controlled temperature (28±0.5°C) and light (10 h dark/14 h light cycle) system with UV-sterilized filtered water (Aquaneering, San Diego, CA). Feeding with rotifers (Rotigrow Nanno, Reed Mariculture, Campbell, CA), brine shrimp (Artemia Brine Shrimp 90% hatch, Aquaneering, San Diego, CA), and flakes (Zebrafish Select Diet, Aquaneering, San Diego, CA) and general assessment of health were performed twice a day. For all assays, randomly selected pairs of adults were placed in 1 liter crossing tanks (Aquaneering, San Diego, CA) in a 1 male:1 female ratio. Embryos from at least five simultaneous crosses were combined. Embryos were then kept in standard Petri dishes with E3 media (0.03% Instant Ocean salt in deionized water) and grown in an incubator at 28±0.5°C, and their health was monitored with a dissecting microscope (Leica, Buffalo Grove, IL). Transgenic lines used for this project were obtained via respective material transfer agreements and included: Tg[*vglut2a*:DsRed] ^52^ from Dr. Hitoshi Okamoto at the RIKEN Brain Science Institute in Japan, Tg[*dlx6a*:GFP] ^51^, and Tg[mpeg1.1:GFP] ^60^ from the Zebrafish International Resource Center. Zebrafish were staged as previously described ^33^. All animal use was approved by the Institutional Animal Care and Use Committee from the Office of Animal Welfare Assurance, University of California, Davis.

### Protein conservation

Coding sequences for the largest transcript for human *SRGAP2* (ENSG00000266028), *SRGAP2C* (ENSG00000171943), mouse *Srgap2* (ENSMUSG00000026425), zebrafish *srgap2* (ENSDARG00000032161), human *SRGAP3* (ENSG00000196220), mouse *Srgap3* (ENSMUSG00000030257), zebrafish *srgap3* (ENSDARG00000060309), human *SRGAP1* (ENSG00000196935), mouse *Srgap1* (ENSMUSG00000020121), zebrafish *srgap1a* (ENSDARG00000007461), and zebrafish *srgap1b* (ENSDARG00000045789) were downloaded from ENSEMBL ^101^. Sequence alignments were performed using the R package *msa* and genetic distances estimated with *seqinr*. Phylogenetic trees were created using the Unweighted Pair Group Method with Arithmetic Mean (UPGMA) with the *hclust* function from the *stats* package. Protein domains were extracted using the UniProtKB/Swiss-Prot database^102^ and conservation estimated with the protein BLAST tool ^103^. Lastly, we used the *Dscript* tool^30^ to predict protein-protein interactions between FBAR domains in human, mouse, and zebrafish SRGAP2 orthologs.

### Protein co-immunoprecipitation

HEK 293T cells were co-transfected with plasmids encoding zebrafish Srgap2-HA and human SRGAP2C-GFP, or zebrafish Srgap2-HA and GFP alone, using the TurboFect™ transfection reagent (Thermo Scientific, R0533) according to manufacturer’s instruction. 24 h after transfection, cells were lysed in 500 µl of Lysis Buffer (20 mM Tris-HCl, pH 8.0, 100 mM KCl, 5 mM MgCl_2_, 0.2 mM EDTA, 10% glycerol, and 0.1% Tween 20) containing 1x protease inhibitor cocktails (Sigma-Aldrich, P8340). The lysates were gently rocked back and forth for 10 min at 4°C and then cleared by centrifugation at 14,000 xg for 5 min at 4°C. 50 µl of the supernatant was saved as the input and the remaining 450 µl was subjected to immunoprecipitation. To capture GFP and GFP fusion proteins, 30 µl of GFP-nanobody conjugated agarose beads—a gift of Henry Ho and prepared as described in ^104^—were washed and blocked with 1 ml of 0.01% bovine serum albumin (BSA) in phosphate-buffered saline (PBS) for 1 h at 4°C before addition of supernatant. The supernatant-beads mix was rocked back and forth for 1 h at 4°C. The beads were then washed with 1 ml of Lysis Buffer three times, 5 min each. The bound proteins were eluted by incubating the beads in 25 µl of 4x Laemmli sample buffer (125 mM Tris-HCl, pH 6.8, 4% sodium dodecyl sulfate, 40% glycerol, 10% 2-mercaptoethanol, and 0.01% Bromophenol blue) at 95°C for 10 min. Proteins in the eluates were then resolved by 10% sodium dodecyl sulfate polyacrylamide gel electrophoresis (SDS-PAGE) and transferred to a polyvinylidene difluoride (PVDF) membrane. After transfer, the PVDF membrane was cut horizontally between 125- and 90-kDa protein markers and blocked in Intercept™ Blocking Buffer (LI-COR, 927-60001) for 1 h at room temperature (RT). The top half was then incubated with the anti-HA antibody (1:10,000 dilution, Invitrogen, 26183) and the bottom half was incubated with the anti-GFP antibody (1:10,000 dilution, Proteintech, 66002-1-lg) in Intercept™ Blocking Buffer for 1 h at RT. After the primary antibody incubation, membranes were washed with Tris-buffered saline (20 mM Tris-HCl, pH 7.6, and 150 mM NaCl) containing 0.1% Tween 20 (TBS-T) three times, 5 min each, and incubated with the IRDye 800RD anti-mouse IgG secondary antibody (1:30,000 dilution, LI-COR, 926-68070) in Intercept™ Blocking Buffer for 1.5 h at RT. Membranes were then washed with TBS-T three times, 5 min each, dried, and imaged using the Odyssey DLx imaging system (LI-COR, Model 9142).

### Baseline expression of *srgap2*

We extract the expression of *srgap2* throughout development from public RNA-seq data that included five biological replicates of pools of 12 embryos at 18 different developmental timepoints ^31^. RNA-seq data from embryonic and adult tissues was retrieved from a recent study ^34^. Raw reads were processed using *fastqc* ^105^, *trimmomatic* ^106^, and *salmon* ^107^ to obtain the transcripts per kilobase million (TPM) values. Validation of *srgap2* temporal expression during development was performed by quantitative PCR (qPCR) at selected timepoints. For this, five NHGRI-1 zebrafish pairs were crossed for each timepoint and three pools of embryos (20 embryos each) collected for whole RNA extraction using the RNeasy kit (Qiagen, Hilden, Germany) with gDNA eliminator columns for DNA removal. The qPCR reactions were prepared following the standard protocol for the Luna kit (New England Biolabs, Ipswich, MA). Oligonucleotide sequences are in Table S2.

### RNA *in situ* hybridization

Whole embryo *in situ* hybridizations were performed as previously described ^108^. Total RNA was extracted from WT zebrafish embryos using Trizol and the riboprobe generated from a pBS-SK-*srgap2* plasmid using a 20 µl *in vitro* transcription reaction containing ∼300 ng of purified plasmid, 2 µl of 10x reaction buffer (New England Biolabs, Ipswich, MA), 2 µl 0.1M DTT, 2 µl of 10x DIG labeling mix 10x DIG labeling mix (Roche, Basel, Switzerland), 0.5 µl of RiboLock RNase inhibitor (Thermo Fisher, Waltham, MA), 0.5 µl of RNA polymerase (T7 or T3), and nuclease-free water. Reactions were incubated at 37°C for 2 h, followed by the addition of 1 µl TURBO DNase (Thermo Fisher, Waltham, MA) and 30 min incubation at 37°C. Reactions were stopped by adding 2 µl of STOP buffer (Promega, Madison, WI). Riboprobe purification was performed with precipitation in 2 µl of 5 M LiCl and 90 µl of 100% ethanol overnight at -80°C. Wild type PTU-treated 24 and 72 hpf embryos were manually dechorionated, fixed in 4% paraformaldehyde in 1x PBS overnight at 4°C, and treated with 10 µg/ml Proteinase K at room temperature for 10 min. The hybridization medium was 65% formamide, 5x SSC, 0.1% Tween 20, 50 µg/ml heparin, 500 µg/ml Type X tRNA, and 9.2 mM citric acid. Embryos were pre-hybridized for 3 h in a 68°C water bath, followed by hybridization with 200 ng of riboprobe in an overnight 68°C water bath. After this, embryos were successively washed at 70°C with hybridization media, 2x SSC, and 0.2x SSC, then with 1x PBS containing 0.1% Tween-20 (1x PBS-Tw) at room temperature. Embryos were incubated for 4 h in blocking solution (2% sheep serum, 2 mg/ml BSA, 1x PBS-Tw), then overnight in blocking solution and 1:5000 diluted anti-DIG antibody (Sigma Aldrich, St. Louis, MO) at 4°C. After incubation, embryos were washed with 1x PBS-Tw and AP buffer (100 mM Tris pH 0.5, 100 mM NaCl, 5 mM MgCl_2_, 0.1% Tween-20) at room temperature right before staining with NBT and BCIP substrates (Roche, Basel, Switzerland) in AP Buffer. Images were obtained using glycerol and a stereomicroscope (M165, Leica, Wetzlar, Germany) with a Leica DFC7000 T digital camera.

### Generation of *srgap2* knockout zebrafish

*srgap2* was disrupted in wild type zebrafish using CRISPR/Cas9 described in previous protocols ^109,110^. The Alt-R system from Integrated DNA Technologies (IDT, Newark, NJ) was used with the following crRNA sequences: GGUCUUGCAGGAGCUGCACACGG (targeting exon 3), CGCUGAUCUGGGCGAAGCGUGGG (targeting exon 4), GAGAGAGUCAGGUGAGCGAGGGG (targeting exon 6), and GUCUCCUGCUAAAUUCCGAAAGG (targeting exon 2). All gRNA sequences were designed using the CRISPRScan tool with the GRCz11/danRer11 genome reference^111^ (sequences found in Table S2). In brief, 2.5 µl of 100 µM crRNA, 2.5 µl of 100 µM tracrRNA (IDT, Newark, NJ), and 5 µl of Nuclease-free Duplex Buffer (IDT, Newark, NJ) were annealed in a program of 5 min at 95°C, a ramp from 95°C to 50°C with a -0.1°C/s change, 10 min at 50°C, and a ramp from 50°C to 4°C with a -1°C/s change. Injection mixes were prepared with 1.3 µl of SpCas9 (20 µM, New England BioLabs, Ipswich, MA), 1.6 µl of annealed crRNA:tracrRNA, 2.5 µl of 4x Injection Buffer (0.2% phenol red, 800 mM KCl, 4 mM MgCl_2_, 4 mM TCEP, 120 mM HEPES, pH 7.0), and 4.6 µl of Nuclease-free water. If several crRNAs were prepared in the same injection mix, equimolar quantities of each crRNA:tracrRNA were included.

We microinjected one-cell-stage zebrafish embryos as described previously ^110^. Briefly, needles were obtained from a micropipette puller (Model P-97, Sutter Instruments) and injections were performed with an air injector calibrated with a microruler (Pneumatic MPPI-2 Pressure Injector). Embryos were collected and ∼1 nl of injection mix injected per embryo. We used two approaches to generate *srgap2* knockouts, one by injecting an injection mix including all 4 gRNAs coupled with SpCas9, and another with an injection mix of the gRNA targeting exon 4 coupled with SpCas9 to create a stable line carrying one specific nonsense mutation. To generate the stable *srgap2* knockout line, we outcrossed our G_0_-injected fish to wild type NHGRI-1 at ∼1.5 months post-fertilization to obtain the G_1_ heterozygous generation, which was further screened by sequencing (EZ-Amplicon sequencing, Azenta, Burlington, MA) a ∼200 bp region that included the gRNA target site (primer sequences in Table S2). Specific alleles were defined using R package *CrispRVariants*^112^. We focused on a 5-bp deletion in exon 4 referred to as *srgap2^tupΔ5^*.

### CRISPR off-target evaluation

Potential off-target sites for the gRNAs were identified from previously generated CIRCLE-seq libraries for each gRNA ^113^, following the standard protocol ^114,115^, and predicted using CRISPRScan ^111^ (Table S3). Injections of each gRNA were performed as described above for subsequent DNA extraction at 5 dpf of injected and non-injected batch-sibling controls, PCR amplification of the top ten off-target sites from each approach (CIRCLE-seq and CRISPRScan), followed by Sanger sequencing (Azenta, Burlington, MA).

### Injection of human mRNA in zebrafish

Temporal expression of *SRGAP2C* mRNA in zebrafish was performed similarly to previously described protocols ^116,117^. The mammalian expression vector pEF-DEST51 containing *SRGAP2C* was used to produce 5’-capped mRNA using the MEGAshortscript T7 transcription kit (Thermo Fisher, Waltham, MA) following the manufacturer’s guidelines with a 3.5 h 56°C incubation with T7 polymerase. mRNA was then purified with the MEGAclear transcription clean-up kit (Thermo Fisher, Waltham, MA), measured using a Qubit (Thermo Fisher, Waltham, MA) and evaluated for integrity by 2% agarose gel electrophoresis. The injection mix contained 100 ng/µl of mRNA, 4x Injection Buffer (0.2% phenol red, 800 mM KCl, 4 mM MgCl_2_, 4 mM TCEP, 120 mM HEPES, pH 7.0), and nuclease-free water. As described above, one-cell stage zebrafish embryos were injected with ∼1 nl of the injection mix and kept at 28°C until needed.

### Morphometric measurements

High-throughput imaging of zebrafish larvae was performed using the VAST BioImager system (Union Biometrica, Holliston, MA) as previously described ^113,118^. In brief, 5 dpf larvae were placed in a rotating 600 µm capillary coupled with a camera, allowing for the automatic acquisition of images from all four sides. Images were automatically processed using FishInspector v1.7 ^119^ to identify and extract morphological shapes, which were then analyzed with the *TableCreator* tool. Images of dead or truncated larvae were discarded. In total, we measured the central line, head area, Euclidean distance between the eyes, and the head-trunk angle across 331 larvae. As no significant differences in measurements of any feature were observed between our controls (uninjected NHGRI-1 wild type larvae, wild type larvae from the stable *srgap2* knockout line, and wild type NHGRI-1 larvae injected with SpCas9 coupled with a scrambled gRNA; all pairwise t-tests p-values > 0.05, complete results in Table S26), we merged these larvae into a single control group.

### Bulk RNA-seq

For stable *srgap2* knockout larvae, a minimum of 3 different *srgap2*^+^/*srgap2^tupΔ5^* x *srgap2*^+^/*srgap2^tupΔ5^*crosses were set embryos were pooled in the batches, and larvae were kept at 28°C until 5 dpf when they were flash frozen and placed in RNA later (Thermo Fisher, Waltham, MA). Tails were then cut off each larva for genotyping via high resolution melt (HRM) curve in a CFX 96 Real-Time System qPCR machine (BioRad). HRM mix included 5 µl DreamTaq DNA polymerase (Thermo Fisher, Waltham, MA), 0.5 µl of each primer at 10 µM, 1 µl of 1x SYBR green (Thermo Fisher, Waltham, MA) and 2 µl of nuclease-free water. In parallel, wild type crosses were set and one-cell stage embryos were injected with human *SRGAP2C* mRNA or the G_0_ knockout. Injections were performed as previously described, using ∼1 nl of the injection mix. For all samples, the heads of five larvae were pooled together and RNA extracted using the RNeasy kit (Qiagen, Hilden, Germany) with gDNA eliminator columns for DNA removal. In total, three samples per group were harvested. Total RNA was then submitted for RNA-seq using poly-A selection and standard library preparation for Illumina sequencing (Genewiz, South Plainfield, NJ).

In a similar manner, 3’-tagged RNA-seq was performed for gene expression evaluations at earlier timepoints. *srgap2* knockouts (stable and pooled), *SRGAP2C*-mRNA injected, and controls were co-injected with SpCas9 and a scrambled gRNA were obtained as previously described. Embryos from each group were collected at 24 hpf (n= 20 larvae per replicate), 48 hpf (n= 10 per sample), and 72 hpf (n= 10 per sample) for flash freezing and incubation in RNAlater (Thermo Fisher, Waltham, MA) at -20°C, completing three replicates per group per timepoint. Once all samples were collected, RNA was extracted from dissected heads using the RNeasy kit (Qiagen, Hilden, Germany). RNA samples were submitted to the UC Davis DNA Technologies Core (Davis, CA) for library preparation and sequencing.

All raw RNA-seq reads were trimmed using *trim-galore* and then mapped to the published zebrafish optimized transcriptome ^120^ using *STAR* ^121^. Gene-level counts were obtained with *HTseq* ^122^. Overall, samples exhibited high correlations in gene counts for both the RNA-seq (mean Spearman ρ= 0.97, range 0.95-0.99) and 3’-tagged RNA-seq (mean Spearman ρ= 0.88, range 0.85-0.93). Differentially expressed genes were obtained with *DESeq2* ^123^ using WT samples from the stable line as controls for the stable knockouts, and injection controls (SpCas9 coupled with a scrambled gRNA) for the G_0_-knockouts and *SRGAP2C*-injected embryos. All enrichment tests of gene groups in specific biological pathways were performed using *clusterProfiler* ^124^ with the background genes including all expressed genes in each dataset (e.g., all genes expressed in microglia cells).

### Single-cell RNA-seq

Embryos were incubated at 28°C. At 3 dpf, the heads of larvae from each group were dissected after euthanasia in cold tricaine (0.025%), pooling 30 heads together per sample (n= three samples per group). Cell dissociation was performed immediately afterward, using previous protocols as reference ^125,126^, with two washes in 1 ml cold 1x PBS on ice and immediate incubation at 28°C for 15 min in a preheated dissociation mix that included 480 µl of 0.25% trypsin-EDTA (Thermo Fisher, Waltham, MA) and 20 µl of collagenase P (100 mg/ml, Sigma-Aldrich, St. Louis, MO). Every 5 min all samples were gently pipetted using a cut P1000 tip. After 15 min, 800 µl stop solution (DMEM with 10% FBS) was added to each sample and immediately centrifuged at 700 g in 4°C for 5 min. The supernatant was discarded and cells were resuspended in cold 1x PBS for another 5 min centrifugation at 700 g in 4°C. After this, the supernatant was discarded and cells were resuspended in 800 µl suspension solution (DMEM with 10% FBS) and filtered through a Flowmi 40 µm cell strainer (Sigma Aldrich, St. Louis, MO) into a low-bind DNA tube (Eppendorf, Hamburg, Germany). Intact cells were counted using a Countess II (Thermo Fisher, Waltham, MA) and cell viability was confirmed to be >65%. Cell fixation and library preparation were then performed with the Parse Biosciences Fixation and Single Cell Whole Transcriptome kit v1.3.0 (Parse Biosciences, Seattle, WA), following the manufacturer’s instructions. A total of 12,500 cells per well were loaded into the barcoding plate and two resulting sub-libraries were sequenced in a NovaSeq 6000 platform.

Raw FASTQ scRNA-seq reads were processed using the Parse Biosciences processing pipeline v0.9.3 and the optimized zebrafish transcriptome ^120^ to obtain the gene x cell matrix files per sample. These matrices were processed into Seurat objects using *Seurat* v4 ^127^. Quality control filtering included feature counts above 200 and below two standard deviations from the mean (5727 features), less than 5% mitochondrial or ribosomal percentages, and doublets removal with *DoubletFinder* ^128^ with a 4% expected doublets for the SPLiT-seq method ^129^. Data for an average of 2391±250 cells per sample were obtained (full sample information in Table S9), which were normalized using *SCTransform* with the top 5,000 variable genes and regressing for mitochondrial and ribosomal percentages. Samples were then integrated using a canonical correlation analysis reduction ^127^ and nearest-neighbor graphs constructed using the first 15 principal components with the *FindNeighbors* function. Hierarchical clustering was performed with the Euclidean distance between principal components embeddings (tree cut at k=40) and cluster marker genes obtained with *PrepSCTFindMarkers* and *FindAllMarkers* using the Wilcoxon test option (parameters: *logfc*.*threshold*= 0.1, *min.pct*= 0.1, *return.thresh*= 0.01, *only.pos*= TRUE), which were further detailed using zebrafish brain atlases ^44,45^ and the ZFIN database ^130^. For the pseudo-bulk analysis, count data was aggregated using *AggregateExpression* and the differential expression test between cell types of different genotypes (e.g., mutant microglia cells vs control microglia cells) performed with the MAST test option ^131^ (parameters: *logfc*.*threshold*= 0.02, *min.pct*= 0.1, *only.pos*= FALSE). Several functions from *scCustomize* ^132^ were used for making plots.

Knockout models exhibited significantly reduced *srgap2* expression (ANOVA genotype effect *p*-value= 3.15x10^-4^, Hom *p*-value= 5.80x10^-4^, G_0_-knockouts *p*-value= 0.011, Table S9), while no reduction was observed in the *SRGAP2C*-injected samples (*SRGAP2C*-humanized *p*-value= 0.992, Table S9), consistent with observations from our quantitative RT-PCR results (Figure S1A). Bulk RNA-seq showed high correlation with single-cell pseudo-bulk gene counts of the same genotype at 3 dpf (average Spearman ρ across genotypes= 0.76±0.03, all *p*-values < 2.2x10^-16^).

### Quantification of neuronal populations

*SRGAP2* models and controls were created as previously described (above) in a Tg[*vglut2*:DsRed] x Tg[*d1x6a*:GFP] background. Embryos were kept at 28°C until 3 dpf, when larvae were anesthetized in tricaine (0.0125%) and embedded in 1% low-melting agarose (n= 6–7 per group). These embryos were imaged using a spinning disk confocal microscope system (Dragonfly, Andor Technology, Belfast, United Kingdom) housed inside an incubator (Okolab, Pozzouli, Italy) with Leica 10x and 20x objectives and an iXon camera (Andor Technology, Belfast, United Kingdom). All imaging was performed using Z-stacking of 10 µm slices starting in the dorsal-most part going ventrally until no fish was detected. Image processing was done using Fiji ^133^ by generating hyperstacks with maximum intensity projections and quantifying all areas either GFP or DsRed positive.

### Motion-tracking activity screen

We performed motion-tracking recordings of 4 dpf larvae using the Zebrabox system with a camera acquisition speed of 30 frames per second (ViewPoint, Montreal, Canada). Larvae were placed in a 96-well plate with 150 µl of E3 media with 0 mM or 2.5 mM pentylenetetrazol (PTZ, #P6500, Sigma-Aldrich, St. Louis, MO) and their movement was recorded for 15 min. A published MATLAB script was used to extract high-speed movement (>28 mm/s) events from data extracted in 1 s bins ^55^ and compared across groups.

### Electrophysiology

Larvae (n= 20–30) at 4 dpf were randomly selected for local field potential (LPF) recordings, as previously described ^55^. Briefly, larvae were exposed to pancuronium (300 µM) and immobilized in 2% low-melting agarose in a vertical slice perfusion chamber (Siskiyou Corporation, #PC-V, Grant Pass, OR). These chambers were then placed on an upright microscope (Olympus BX-51W, Lausanne, Switzerland) and monitored with a Zeiss Axiocam digital camera. 15 min LFP recordings were obtained by placing a single-glass microelectrode (WPI glass #TW150 F-3) with a ∼1 µm tip diameter in the optic tectum under visual guidance. The voltage signals were filtered at 1 kHz and digitized at 10 kHz using Digidata 1320 A/D interface (Molecular Devices, San Jose, CA). All recordings were coded and scored independently by three researchers using Clampfit software (Molecular Devices, San Jose, CA) to obtain the final LFP score per group.

### Histology and immunostaining

We evaluated the general morphology of the eye in 5 dpf larvae and performed immunohistochemistry using anti-Pax6 antibodies (Thermo Fisher, Waltham, MA) to label the amacrine and retinal ganglion cells in the eyes. In brief, 10 µm sections for each group were collected using a cryostat microtome (Leica, Wetzlar, Germany) and placed on slides at -80°C. Slides were then brought to room temperature and washed with 1 ml 1x PBS for 5 min, followed by incubation with blocking buffer (4% milk/TST buffer) for 1 h. Then, the blocking buffer was removed, and slides were incubated with anti-Pax6 antibodies in blocking buffer overnight at 4°C. Incubation with a secondary anti-mouse antibody (Thermo Fisher, Waltham, MA) was performed for 1 h after a wash with fresh blocking buffer. Images were obtained using a confocal microscope (Olympus, Lausanne, Switzerland). Additionally, cryosections (10 µm) from each group were stained with hematoxylin and eosin (H&E) and mounted in Permount.

### Visual-motor response assays

We performed visual-motor response tests on 5 dpf larvae in a 96-well plate with 150 µl E3 media per well (n= 24 per group). Using the Zebrabox system (ViewPoint, Montreal, Canada), we exposed larvae to a protocol consisting of 10 min dark adaptation followed by bright light (100 lumens) and recorded their movement responses. Movement data were exported in 1 s bins for comparisons across groups in the 20 s prior and post dark-to-light change. Additionally, we performed optomotor response (OMR) tests following a protocol that uses a monitor to display a video with 30 s periods of contrasting stripes moving at 1.04 rad/s separated by 20s intervals ^73^. We placed 4 larvae per group in a standard Petri dish and exposed them to 5 cycles of the recording, with 3 replicates per group (n= 12 larval measurements per group). In separate experiments, video recordings were paused during every cycle, after exactly 10 s (halfway through the video) and the number of larvae with rostral ends oriented in the direction of the moving stripes was recorded, giving the “OMR positive” response. The quantification was performed blind to genotype.

### Microglia morphology and abundance

One-cell stage larvae from a Tg[*mpeg1.1*:GFP] cross were microinjected as described above to generate *srgap2* G_0_-knockouts, *SRGAP2C*-injected, and scrambled gRNA-injected controls. At 3 and 7 dpf, larvae were anesthetized with MS-222 (0.175 mg/ml in E3 media), embedded in 1% low-melt agar, and immediately imaged with a spinning disk confocal microscope system fitted with a 63X lens (Dragonfly, Andor Technology, Belfast, United Kingdom) as described above. Sphericity was obtained as described ^62,134^ using the Imaris software (Bitplane, Switzerland) and creating 3D surface reconstructions per cell. Parameters were consistent across samples, including a smooth selection of 0.191µm and thresholding of absolute intensity. A total of one to five microglial cells were imaged from three to four larvae per genotype per timepoint. Microglial cells were quantified according to an established protocol ^59,135^. 3dpf larvae were incubated in E3 media containing 2.5 μg/ml neutral red at 28.5°C for 3 hr, followed by two water changes and imaged immediately after using a stereoscope (M165, Leica, Wetzlar, Germany) with a Leica DFC7000 T digital camera.

### Human and non-human primates scRNA-seq

scRNA-seq data from human retinal organoids ^68^ (43,857 cells), human donors ^70^ (183,808 cells), macaque retinal organoids ^69^ (19,894 cells), macaque donors ^70^ (165,681 cells), and prefrontal cortex data from humans and non-human primates^66^ (171,997 human cells, 158,099 chimpanzee cells, 131,032 macaque cells, 149,468 marmoset cells) were downloaded as preprocessed objects. Retinal datasets were integrated using the LIGER method for cross-species analyses ^136^ followed by joint matrix factorization with *optimizeALS* using a lambda of 5, a convergence threshold of 1x10^-10^, and a k of 30. Differentially expressed genes were obtained with *getFactorMarkers*, using the human data as reference. Enrichment of genes in biological pathways was performed using *clusterProfiler* ^124^. For the prefrontal cortex data ^66^, we obtained differentially expressed genes with the *FindMarkers* function from Seurat v4.0 ^127^ using the Wilcoxon test option. Microglial cells defined in the prefrontal cortex ^66^ and the middle temporal gyrus ^137^ were gathered totaling 30,918 cells (prefrontal cortex: human= 7,556, chimpanzee= 5,748, macaque= 8,058, marmoset= 4,626; middle temporal gyrus: human= 1,263, chimpanzee= 252, macaque= 942, marmoset= 2473) and their expression aggregated using *AggregateExpression* from Seurat ^127^, grouping by organism to obtain a gene by organism pseudo count table. Differential gene expression between species was then performed with *DESeq2* ^123^ and overrepresentation tests in GO terms with DAVID ^138^.

### Statistical analysis

All statistical analyses were performed in R version 4.0.2, and all scripts are available in the GitHub repository https://github.com/mydennislab/public_data/ (zenodo pending). Comparisons between groups were performed using two-tailed Student’s t-tests, Mann-Whitney U-tests, Analysis of Variance (ANOVA) or nonparametric Dunn’s tests, depending on the normality of the data assessed using the Shapiro-Wilk test. All analyses across experimental batches included *batch* as a factor in the model to control for biases caused by inter-batch differences. Fisher’s exact tests were used for testing significant overlaps between gene lists. All mean values reported include their standard deviation unless otherwise noted. Significance thresholds were defined with an alpha of 0.05 and the proper corrections for multiple comparisons defined in the text. All gene ontology enrichment tests were performed using solely the expressed genes as the background gene list.

## Data availability

GEO numbers of deposited data pending: bulk RNA-seq, scRNA-seq

## Acknowledgements

We thank Kyle Burbach, Daisy Castillo, Aarthi Sekar, Dr. Alexandra Colón-Rodríguez, and Eva Ferino for their support in helping to generate and maintain the *srgap2* stable knockout line. We also thank Dr. Colin Shew, Jennielee Mia, Xueer Jiang, and Ingrid Brust-Mascher for technical support in computational and imaging analyses. We thank Drs. Bruce Draper, Kristen Kwan, Heather Mefford, Anna La Torre, Ala Moshiri, Nick Marsh-Armstrong, and Paul FitzGerald for many fruitful discussions, ideas, reagents, and advice. We thank the Cell Biology and Human Anatomy Department at UC Davis School of Medicine for the support to use multiple pieces of imaging equipment. We also thank Dr. Elizabeth Haswell, as well as Aidan Baraban and Gabriana La for significant edits to the manuscript. This work was supported, in part, by the U.S. National Institutes of Health (NIH) grants from the Office of the Director and National Institute of Mental Health (DP2MH119424 and R01MH132818 to M.Y.D.), UC Davis MIND Institute Intellectual and Developmental Disabilities Research Center pilot grant (U54 HD079125 to M.Y.D.), NIH National Institute of General Medical Sciences (NIGMS) (R01GM144435 to L.-E.J.), NIH National Institute of Neurological Disorders and Stroke (R01NS096976 and R01NS103139 to S.C.B, and R01NS109176 to S.S.). NKH is supported by an NIH NIGMS UC Davis eMCDB T32 (T32GM153586), and a UC Davis Graduate Research Award (J.M.U-S.).

