## Supplementary Figures for "Zebrafish models of human-duplicated *SRGAP2* reveal novel functions in microglia and visual system development"

**A**

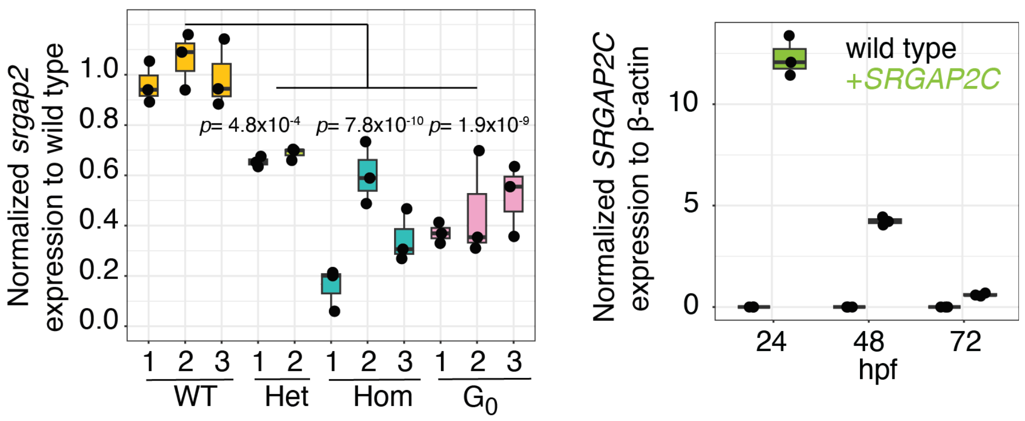


**B**

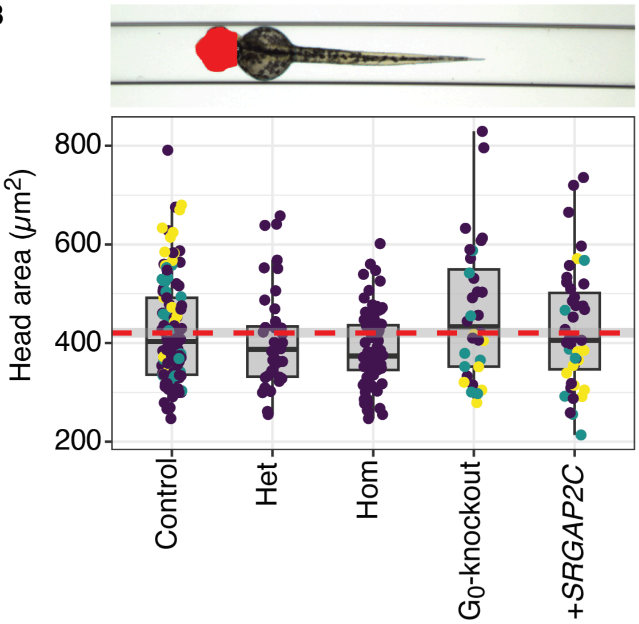


**Figure S1. *SRGAP2* zebrafish models (Related to Figure 2). (A)** Expression of *srgap2* in knockouts from the stable line (Hom, Het, wildt type) and $G_0$-knockouts, and expression of *SRGAP2C* in uninjected (wildt type) and *SRGAP2C*-injected (+*SRGAP2C*) larvae at 5 dpf. Each dot represents a biological replicate (pool of n=20 larvae). Dunnett's test FDR-adjusted *p*-values: Het= $4.8\times10^{-4}$, Hom= $7.8\times10^{-10}$, $G_0$-knockout= $1.9\times10^{-9}$. (**B)** Measurements of head area (ANOVA: $F_{(4, 315)}$= 3.90, genotype effects *p*-value= 0.00416, FDR-adjusted *p*-values Het= 0.987, Hom= 0.271, Pooled= 0.304, *SRGAP2C*= 0.987). Dots represent an imaged larva with the color indicating the imaging plate (a co-variable included in the statistical analyses). The red dotted line corresponds to the mean value for the control group, and the red asterisks on top indicate an FDR-adjusted Dunnett's *p*-value < 0.05. A representative image is included on the top of the plot.

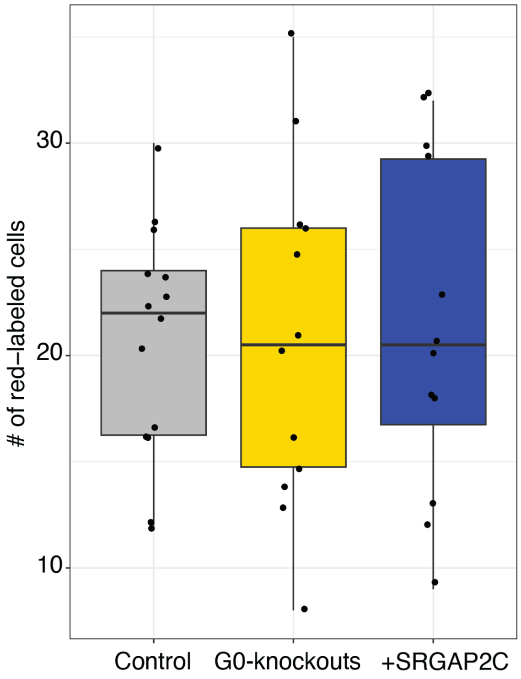


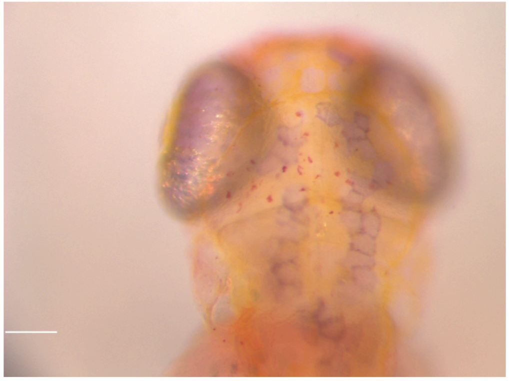

**Figure S2. Microglial cell abundance in *SRGAP2* mutants (Related to Figure 4).** A neutral red assay was used to quantify microglial/myeloid cells in *srgap2* knockout (n=12), *SRGAP2C*-injected (n=14), and scrambled gRNA-injected control larvae (n=12). Each dot in the graph represents an imaged larvae. A representative image is shown below the graph with the red-positive cells.

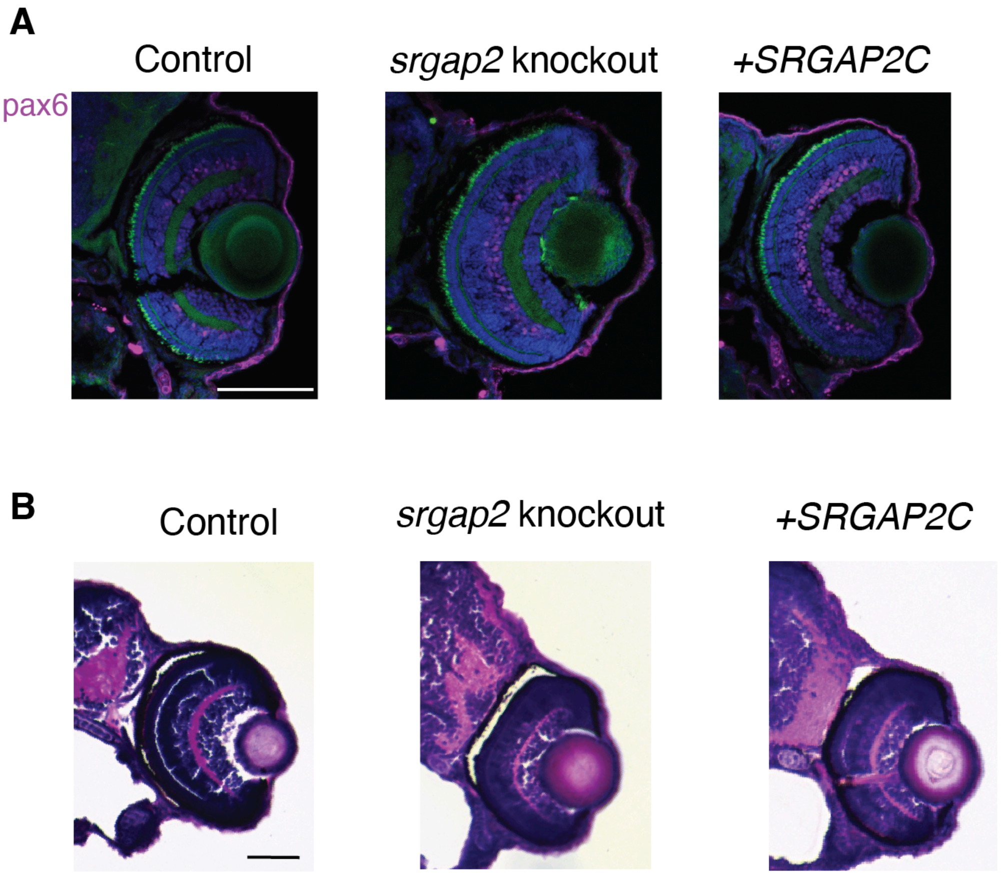


**Figure S3. Gross eye morphology of *SRGAP2* mutants (Related to Figure 5).** (**A**) Immunohistochemistry of 5 dpf SpCas9-scrambled gRNA controls, stable knockouts, and *SRGAP2C*-injected larvae using anti-Pax6 antibodies to label amacrine and retinal ganglion cells. All retinal tissue layers were identified with no obvious differences across groups. (**B)** Hematoxylin and eosin staining of 3 dpf SpCas9-scrambled gRNA controls, stable knockouts, and *SRGAP2C*-injected larvae, evidencing no gross differences in retinal structure organization across groups.
