## Supplementary Notes for "Zebrafish models of human-duplicated *SRGAP2* reveal novel functions in microglia and visual system development"

### Supplementary Note

**Note S1. Transcriptomic correlation of *SRGAP2* zebrafish models (related to Figure 2).** In our transcriptome data of 5 dpf larvae, we noted a subgroup of commonly upregulated genes with relative higher expression in the *SRGAP2C*-injected versus the $G_0$-knockout larvae that were enriched in visual system development terms, including multiple crystallins and small heat-shock proteins (e.g., *cryaa*, *cryba1b*), in line with recent published work [1] in which orthologs of these same genes were also increased in the retina of *Srgap2*$^{+/-}$ mice. We also observed high concordance in the expression pattern of $G_0$-knockouts and stable Hom larvae (Spearman ρ for the DEGs=0.39, *p*-value= 0.010), despite the smaller number of DEGs observed in the Hom (n=57, Table S6), giving us further confidence in the observed patterns.

**Note S2. *SRGAP2* paralog expression profiles in great ape post-mortem samples (related to Figure 4).** Leveraging the detailed classification of microglial cell types performed in the original study [2] in which they identified three microglial subtypes—one subtype that is common between all species (Micro, main marker gene: *P2RY12*), one subtype found only in human and chimpanzees termed "Hominidae microglia" (hoMicro, main marker gene: *GDLN*), and a third subtype solely found in human samples termed "human microglia" (huMicro, main marker genes: *CCL4*, *CCL3*) we were able to assess the level of expression of *SRGAP2* and *SRGAP2C* amongst species (Figure 4C). Importantly, these microglial subtypes that were restricted to some species were found in low abundances relative to total microglial cells (hoMicro: human 5.8%, chimpanzee 1.2%, huMicro: 4.1%). Overall, all microglial subtypes retained high expression of *SRGAP2* across species, with huMicro subtype showing slightly higher average levels of expression compared to the common Micro subtype.

**Note S3. Comparison of human and rhesus macaque retinal transcriptomic profiles (related to Figure 5).** Motivated to further investigate the potential role of *SRGAP2* in the development of human retinal features, we leveraged public single-cell RNA-seq data from human and macaque retinal organoids [3,4] to identify genes with increased expression in humans. We integrated 63,751 cells (43,857 human and 19,894 macaque) and found 777 and 767 genes with increased expression in humans and macaques, respectively (Figure 5C, Table S21). Strikingly, genes with increased expression in human retinal cells were overrepresented in axogenesis-related and light perception GO terms, in line with our results and the known functions of *SRGAP2* (Table S22). Genes with increased expression in macaque cells were enriched in functions for salt/calcium response and miRNAs regulation. To validate these results, we performed the same analysis using 418,808 cells (183,808 human and 165,681 macaque) from post-mortem donors [5,6] and obtained the same overrepresentation of genes with increased expression in humans in axogenesis-related GO terms (Tables S21 and S23). Moreover, we intersected the 777 genes with increased expression in human retinal organoids with the commonly upregulated genes in the RGC and RPE our *SRGAP2* mutants (n= 487 genes) and observed a significant overlap (Fisher's test odds ratio= 6.23, *p*-value< $2.2 \times 10^{-16}$, 77 overlapping genes, full list in Table S24), evidencing that our observed signals using zebrafish models represent real mechanisms. In all, these results highlight the importance of detailing the role of *SRGAP2* in the development of human features of the retina, which is yet unexplored.
